# Indoor environmental conditions shape the microbial landscape of food production facilities across space and time

**DOI:** 10.1101/2025.08.07.669043

**Authors:** Nicholas A. Bokulich, Eve Beauchemin, Chad Masarweh, Irnayuli Sitepu, Karen Kalanetra, Roger Boulton, David A. Mills, Kyria Boundy-Mills

**Affiliations:** Department of Health Sciences and Technology, ETH Zurich, Zurich, Switzerland; Department of Food Science and Technology, University of California, Davis, Davis, CA, USA; Department of Viticulture and Enology, University of California, Davis, Davis, CA, USA

## Abstract

The microbial communities inhabiting food production environments are distinguished from those of other built environments in their capacity to influence food quality and safety, impacting consumer health. However, how indoor environmental conditions drive the bacterial and fungal communities of food production facilities remains largely unknown. In this study of five commercial food production facilities, we employed remote wireless sensors paired with marker-gene amplicon sequencing (bacterial 16S rRNA genes and fungal ITS) of processing equipment and non-processing built environment surfaces (N=2,329) to profile spatial and longitudinal changes in bacterial and fungal communities, and their association with indoor climate. We identify multiple associations between indoor environmental conditions and microbial community structure, demonstrating the role of the indoor environment in shaping microbial communities on food processing and non-processing surfaces. This suggests that indoor climate could be manipulated to rationally modify surface communities, with the aim of enhancing food quality and safety.

## Introduction

Food supply chains in industrialized countries are highly contained, with the vast majority of the human diet being processed, packaged, stored, and transported within indoor environments prior to consumption. As with other indoor environments (1), food processing facilities present a unique set of conditions that influence microbial community structure and activities, including the presence of building materials, substrates, and physicochemical conditions that significantly differ from natural ecosystems. The impact of these conditions on the adaptation and ecological assembly of microbiomes in food processing ecosystems is largely unexplored. The microbial ecosystems of food production play critical roles in food quality, spoilage, and safety (2–4). Consequently, surveillance of food processing facilities throughout the food supply chain is paramount to understanding the role environmental microbiota play in shaping holistic product qualities.

Fermented-food processing facilities are unique among indoor environments in that microbial activity is an inherent feature of the production process, impacting product quality and consumer acceptance. Microbial activity is responsible for both beneficial quality characteristics as well as spoilage and quality deterioration in various fermented foods (2–4). However, the ecology of these organisms within food-processing environments is less understood, in particular how indoor environmental conditions shape the distribution of microbial species within food facilities. High-throughput marker-gene sequencing methods have enhanced the ability to study complex microbial communities within fermented foods and food facilities (5–15), offering an opportunity for exploring the connection between indoor environmental conditions and microbial community compositions in food facilities (16). In this work we focus on food facilities producing wine and dairy products, as two economically and dietarily important food groups for which microbial activity crucially shapes quality, safety, and spoilage.

In winery ecosystems, the origins of microbes in wine fermentations (other than those that are intentionally inoculated) are poorly understood and frequently assumed to be from grapes (17). However, many of the primary microbes involved both positively and negatively in wine fermentations are only detected as minor populations on the surface of healthy grapes, if at all (17, 18). An important source for the transfer of these microbes between fermentations is the winery environment itself (19, 20). The distribution of non-*Saccharomyces* yeasts (19, 21)(22), lactic acid bacteria (23), and molds (24, 25) has also been investigated in wineries using culture-based techniques. High-throughput marker-gene sequencing of bacterial and fungal communities reveals the rich microbial ecosystems and potential reservoirs for microbial contaminants within winery environments (12, 20). Importantly, the winery microbial ecosystem fluctuates seasonally, apparently following grape harvest and processing schedules, and thus in response to the movement and conversion of fermentation substrates within the facility (12). These fluctuations may also reflect changes in ambient conditions following seasonal weather conditions. However, the indoor conditions that drive microbial fluxes within winery environments are unknown and have not been considered in previous studies. Given the evidence for the interface between winery surface microbiota and wine fermentations, identifying these conditions may help manage microbial landscapes and prevent the establishment of detrimental microbial reservoirs within winery environments, leading to improved wine quality.

The microbial ecosystem of cheesemaking facilities is different from in wineries, given differences in substrates, processing techniques, and indoor conditions. During cheese production, the product encounters many equipment surfaces on its journey from milk to curd to cheese, all acting as potential vectors for microbes. Hence, the processing environment serves as an important reservoir for bidirectional microbial transfer between fermentations (5, 8, 26). Biofilms of psychrotrophic bacteria (27, 28) and non-starter lactic acid bacteria (29, 30) can form on equipment surfaces, acting as a source of contamination in successive batches of cheese. Wooden processing surfaces, including aging boards (31, 32), milk vats (7, 33, 34), and brining equipment (6) are also rich sources of microbes that are important for cheese acidification and ripening. Microbial transfer from processing surfaces to cheese fermentations can influence quality characteristics (33, 34), and hence the unique microbial communities present in different cheesemakers could be important contributors to the characteristics of regional and artisanal cheeses (5). However, the conditions influencing the formation and transfer of microbial communities within dairies are poorly understood.

In the current study, we employ high-throughput marker gene sequencing of bacteria (16S rRNA gene sequencing) and fungi (ITS domain sequencing) to examine temporal trends in the spatial distribution of microbial communities within 3 wineries and 2 cheese makers over the course of up to 1 year. We predicted that both sample type (as we have shown previously (5)) and indoor environmental conditions would impact the microbial communities inhabiting equipment and environmental surfaces. Remote wireless sensors were placed throughout these facilities for continuous monitoring of indoor environmental conditions. We integrated these environmental and microbial data to demonstrate that the microbial ecosystems inhabiting food facilities are responsive to indoor conditions.

## Results

### Facility and sample type are the primary variables associated with microbial community composition

Microbiome sequencing was performed on surface swabs, raw materials, and fermented food products collected in 5 fermented food production facilities to profile changes in bacterial and fungal community compositions across space and time. We have previously found that sample type is associated with microbial community compositions in various food production facilities (5, 9, 10, 12), and sought to recapitulate this finding across a range of food producers in the current study. We define “sample type” to encompass both different materials (soil, raw materials used in food production, fermented food products, waste streams) and exposure to different production practices (e.g., processing equipment used in food production vs. building surfaces that do not directly contact foods) (Tables S1, S2). We also categorized samples by “room type”, indicating the particular function of different production areas (e.g., areas for raw material handling vs. main production areas), and to examine whether spatial partitions could impact microbial community compositions.

To quantify spatial variation in the diversity of microbiomes on these different surfaces, we used multiple beta-diversity metrics to quantify the similarity between microbial communities observed in each sample: abundance-weighted and unweighted UniFrac distances (35), which measure the phylogenetic similarity between microbial communities; Bray-Curtis dissimilarity (36), a non-phylogenetic, abundance-weighted method; and Jaccard distance (37), a non-phylogenetic, unweighted method. For the Bray-Curtis and Jaccard metrics, our input consisted either of the traditional ASVs or k-mer frequencies (see Methods), the latter of which can provide information on the relatedness between ASVs, since more closely phylogenetically related organisms typically share more k-mers in common.(38)

We used PERMANOVA tests (39) to determine whether these microbial diversity metrics partition by sample type, room type, facility, and multiple environmental measurements (Tables 1-4); in other words, whether these variables are associated with differences in the microbial community compositions of each sample. For the ASV-based metrics, sample type was found to explain the greatest percent of variation in both bacterial and fungal communities in wineries across all metrics (R^2^ min = 0.07, max = 0.15, P < 0.05, Tables 2 and 4); it was also a major factor in creameries, though facility was the primary driver in both bacterial and fungal communities in creameries (R^2^ min = 0.06, max = 0.38, P < 0.05, Tables 1 and 3), consistent with our prior discovery of facility-specific “house microbiota” in these two cheese makers (5). Room type was significantly associated with bacterial and fungal compositions in both wineries and creameries according to most diversity metrics, but contributed a smaller amount of variation (R^2^ min = 0.02, max = 0.08, P < 0.05, Tables 1-4). The k-mer-based metrics were largely concordant with the ASV-based metrics, with the exception that sample type typically explained more variation in creameries than did facility (Table S3), a reversal of the trend seen with ASV-based non-phylogenetically-aware metrics (Tables 1 and 3). These results suggest that, although there are significant differences in the composition of sequence variants observed between these creameries, these represent phylogenetically related groups (e.g., different strains of the same species inhabit the different facilities), whereas different sample types recruit phylogenetically distinct clades. This result is expected, as the general substrates, environmental conditions, and management practices in creameries likely select for closely related (similar) microorganisms with similar abilities to survive in this environment.

**Table 1.** PERMANOVA analysis of creamery 16S rRNA data based on four distance/dissimilarity metrics.

|  |  | Bray Curtis |  | Jaccard |  | Weighted UniFrac |  | Unweighted UniFrac |  |
| --- | --- | --- | --- | --- | --- | --- | --- | --- | --- |
| Variable | Df | R<sup>2</sup> | P-value | R<sup>2</sup> | P-value | R<sup>2</sup> | P-value | R<sup>2</sup> | P-value |
| Facility | 1 | 0.12 | **0.001** | 0.08 | **0.001** | 0.13 | **0.001** | 0.06 | **0.001** |
| Room Type | 4 | 0.07 | **0.002** | 0.06 | **0.002** | 0.08 | **0.003** | 0.06 | **0.013** |
| Sample Type | 6 | 0.10 | **0.001** | 0.08 | **0.002** | 0.13 | **0.001** | 0.10 | **0.002** |
| Daily Mean Temperature | 1 | 0.01 | 0.139 | 0.01 | 0.082 | 0.02 | 0.086 | 0.01 | 0.118 |
| Daily Mean CO2 | 1 | 0.02 | **0.028** | 0.02 | **0.002** | 0.02 | 0.054 | 0.01 | 0.108 |
| Daily Mean Humidity | 1 | 0.02 | 0.053 | 0.02 | **0.010** | 0.01 | 0.179 | 0.01 | 0.109 |
| Weekly CV CO2 | 1 | 0.02 | **0.018** | 0.01 | 0.074 | 0.02 | **0.015** | 0.01 | 0.205 |
| Weekly CV Humidity | 1 | 0.01 | 0.305 | 0.01 | 0.423 | 0.01 | 0.433 | 0.01 | 0.464 |
| Weekly CV Temperature | 1 | 0.01 | 0.135 | 0.01 | 0.174 | 0.01 | 0.192 | 0.01 | 0.453 |
| Residuals | 73 | 0.62 |  | 0.69 |  | 0.57 |  | 0.71 |  |

\* abbreviations: CV = coefficient of variation; Df = degrees of freedom. Significant P-values (P < 0.05) shown in boldface text.

**Table 2.** PERMANOVA analysis of winery 16S rRNA data based on four distance/dissimilarity metrics.

|  |  | Bray Curtis |  | Jaccard |  | Weighted UniFrac |  | Unweighted UniFrac |  |
| --- | --- | --- | --- | --- | --- | --- | --- | --- | --- |
| Variable | Df | R<sup>2</sup> | P-value | R<sup>2</sup> | P-value | R<sup>2</sup> | P-value | R<sup>2</sup> | P-value |
| Facility | 2 | 0.06 | **0.001** | 0.04 | **0.001** | 0.05 | **0.001** | 0.03 | **0.001** |
| Room Type | 4 | 0.02 | **0.002** | 0.02 | **0.001** | 0.02 | 0.272 | 0.02 | **0.018** |
| Sample Type | 7 | 0.10 | **0.001** | 0.07 | **0.001** | 0.15 | **0.001** | 0.08 | **0.001** |
| Daily Mean Temperature | 1 | 0.01 | **0.024** | 0.01 | **0.001** | 0.01 | **0.006** | 0.01 | **0.003** |
| Daily Mean CO2 | 1 | 0.01 | 0.064 | 0.01 | **0.036** | 0.01 | **0.032** | 0.01 | **0.009** |
| Daily Mean Humidity | 1 | 0.01 | **0.001** | 0.01 | **0.001** | 0.01 | **0.022** | 0.01 | **0.001** |
| Weekly CV CO2 | 1 | 0.01 | **0.008** | 0.01 | **0.004** | 0.01 | **0.040** | 0.01 | **0.024** |
| Weekly CV Humidity | 1 | 0.01 | **0.049** | 0.01 | **0.037** | 0.01 | 0.198 | 0.01 | 0.208 |
| Weekly CV Temperature | 1 | 0.01 | 0.129 | 0.01 | 0.099 | 0.00 | 0.519 | 0.01 | 0.214 |
| Residuals | 178 | 0.78 |  | 0.83 |  | 0.74 |  | 0.82 |  |

\* abbreviations: CV = coefficient of variation; Df = degrees of freedom. Significant P-values ( $P < 0.05$ ) shown in boldface text.

**Table 3.** PERMANOVA analysis of creamery ITS data based on Bray Curtis dissimilarity and Jaccard distance.

|  |  | Bray Curtis |  | Jaccard |  |
| --- | --- | --- | --- | --- | --- |
| Variable | Df | R<sup>2</sup> | P-value | R<sup>2</sup> | P-value |
| Facility | 1 | 0.38 | **0.001** | 0.11 | **0.001** |
| Room Type | 4 | 0.06 | **0.008** | 0.04 | 0.254 |
| Sample Type | 6 | 0.10 | **0.001** | 0.09 | **0.002** |
| Daily Mean Temperature | 1 | 0.02 | **0.025** | 0.01 | 0.117 |
| Daily Mean CO2 | 1 | 0.01 | 0.081 | 0.01 | 0.583 |
| Daily Mean Humidity | 1 | 0.01 | 0.243 | 0.01 | 0.264 |
| Weekly CV CO2 | 1 | 0.01 | 0.284 | 0.01 | 0.091 |
| Weekly CV Humidity | 1 | 0.01 | 0.360 | 0.01 | 0.545 |
| Weekly CV Temperature | 1 | 0.01 | 0.107 | 0.01 | 0.617 |
| Residuals | 70 | 0.40 |  | 0.69 |  |

\* abbreviations: CV = coefficient of variation; Df = degrees of freedom. Significant P-values ( $P < 0.05$ ) shown in boldface text.

**Table 4.** PERMANOVA analysis of winery ITS data based on Bray Curtis dissimilarity and Jaccard distance.

|  |  | Bray Curtis |  | Jaccard |  |
| --- | --- | --- | --- | --- | --- |
| Variable | Df | R<sup>2</sup> | P-value | R<sup>2</sup> | P-value |
| Facility | 2 | 0.07 | **0.001** | 0.05 | **0.001** |
| Room Type | 4 | 0.04 | **0.001** | 0.03 | **0.001** |
| Sample Type | 6 | 0.10 | **0.001** | 0.07 | **0.001** |
| Daily Mean Temperature | 1 | 0.02 | **0.001** | 0.01 | **0.001** |
| Daily Mean CO2 | 1 | 0.01 | **0.008** | 0.01 | **0.001** |
| Daily Mean Humidity | 1 | 0.01 | **0.001** | 0.01 | **0.001** |
| Weekly CV CO2 | 1 | 0.01 | 0.051 | 0.01 | **0.013** |
| Weekly CV Humidity | 1 | 0.01 | **0.006** | 0.01 | **0.001** |
| Weekly CV Temperature | 1 | 0.01 | **0.036** | 0.00 | 0.156 |
| Residuals | 199 | 0.73 |  | 0.80 |  |

\* abbreviations: CV = coefficient of variation; Df = degrees of freedom. Significant P-values ( $P < 0.05$ ) shown in boldface text.

Principal coordinate analysis (PCoA) allows us to visualize sample dissimilarities in two-dimensional space. Samples roughly clustered by sample type across multiple facilities and with multiple diversity metrics (Figure 1, Figure S1), supporting indications from PERMANOVA that similar sample, facility, and room types exert similar impacts on microbial communities in each facility. In both creameries and wineries, and for both bacterial and fungal communities, fermented food products and raw materials cluster away from environmental surfaces, indicating that microbial communities in food materials and fermented products are differentiable from those in the surrounding environment. However, the relatively weak separation between food materials (raw materials and fermented food products) and the environment indicate a large degree of microbial sharing; this is particularly prominent with Jaccard distance (the fraction of features not shared between each pair of samples), which exhibits very low PERMANOVA R^2^ values and principal coordinate weights in axes 1-3, both of which relate to the percent of variation explained. In general, processing equipment centroids appear in an intermediate position between raw material and/or fermented food product centroids and building surface swabs (i.e., non-production surfaces), indicating a higher degree of community similarity and microbial sharing between foods and processing surfaces, compared to non-processing surfaces (Figure 1). Pairwise PERMANOVA tests indicate that the fungal and bacterial communities in each of these sample types (raw materials, fermented food products, processing equipment, and building surfaces) are significantly different (P < 0.05), in both creameries and wineries, though raw materials and fermented food products were not distinguishable in a minority of tests – likely because the category “fermented food product” includes active fermentations (Tables S4 and S5). The relative similarities between raw materials, fermented food products, and processing surfaces are qualitatively apparent in the bar plots presented in Figure 2, which represent the average relative taxonomic composition of bacterial and fungal communities in each sample type. For instance, raw materials, fermented food products, and processing surfaces seem to share higher levels of *Lactococcus* in creameries and slightly higher levels of *Saccharomyces* and *Acetobacteraceae* in wineries (Figure 2).

**Figure 1.**
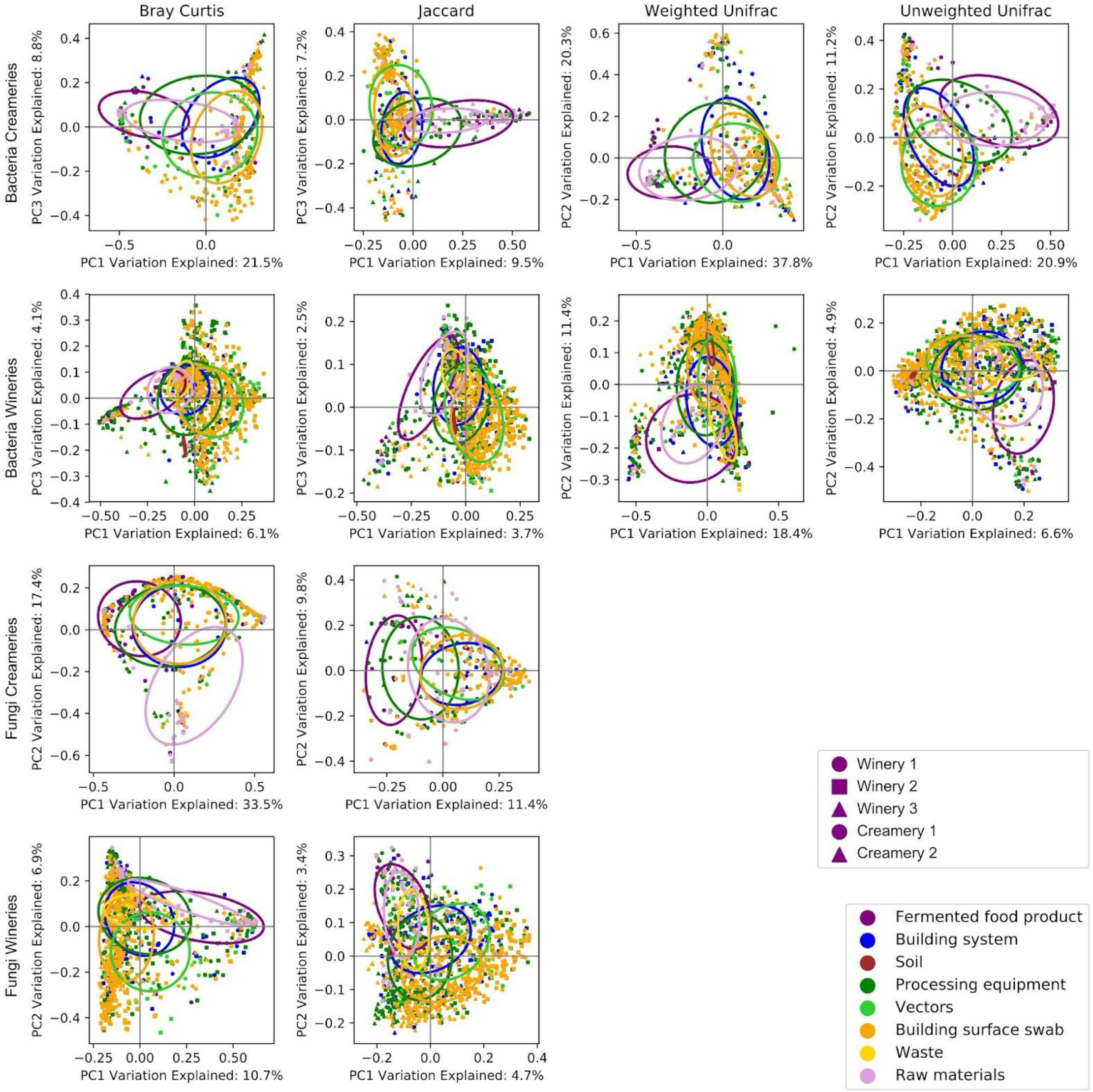
Sample type is the principal driver of bacterial and fungal community composition in creameries and wineries. Principal coordinate analysis of winery and creamery samples, representing both bacterial 16S rRNA (V4) and fungal ITS data evaluated using four different distance metrics. Samples (points) are color-coded by sample type and shape represents different facilities. Ellipses represent one standard deviation from the centroid of each sample type.

**Figure 2.**
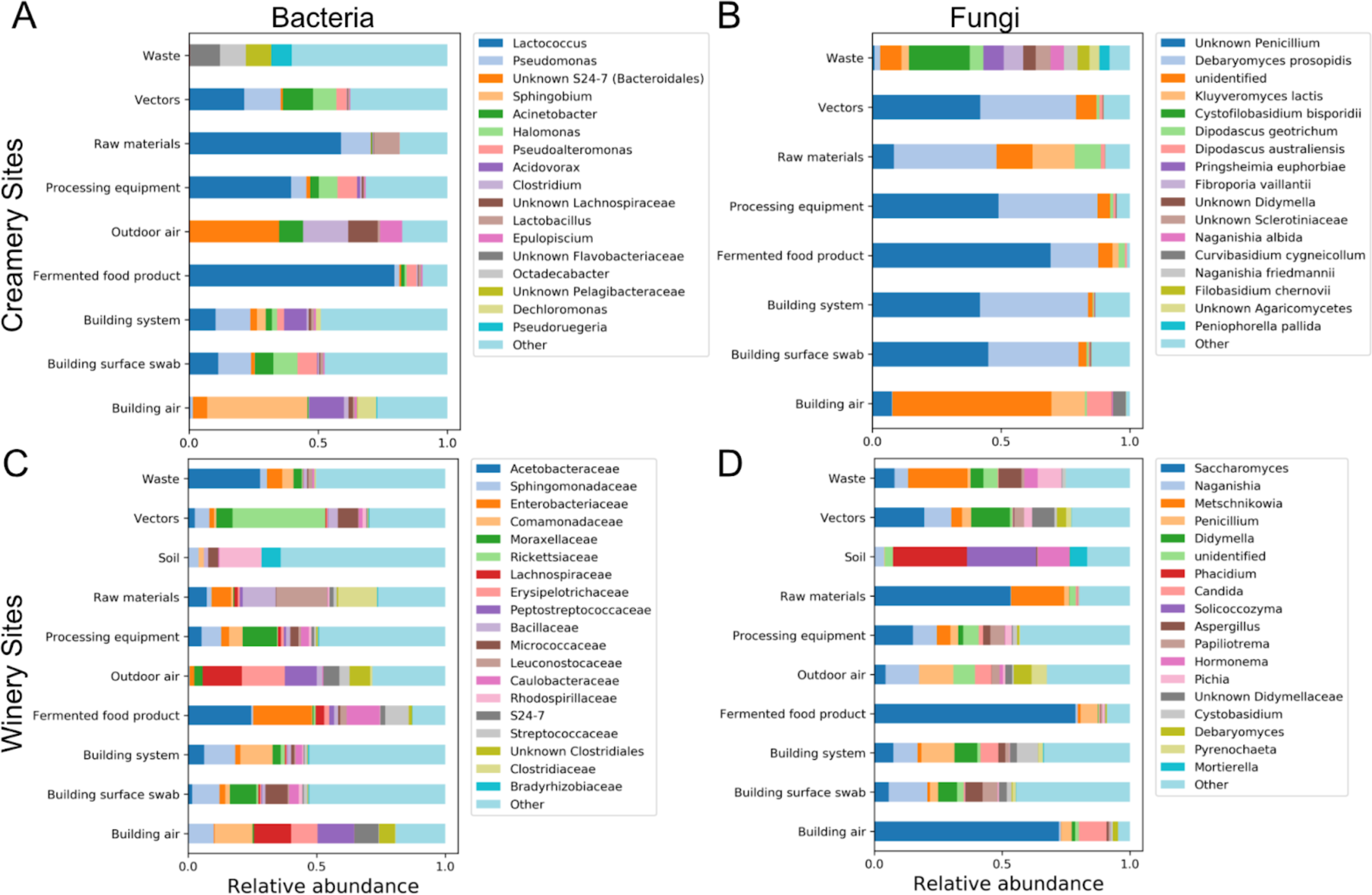
Mean bacterial (left) and fungal community compositions (right) for each sample type. Microbial abundances are averaged across all facilities and all samples of each type. Only the most abundant taxa are shown: these are taxa that were detected at a mean relative abundance (RA) ≥ X in at least one sample type, where X equals the threshold specified. Thresholds were chosen programmatically to limit to the top most abundant taxa. All remaining taxa are aggregated into “Other”. **A,** Genus-level bacterial community abundances in creamery samples, threshold RA ≥ 0.07. **B,** Species-level fungal community abundances in creamery samples, threshold RA ≥ 0.03. **C,** Family-level bacterial community abundances in winery samples, threshold RA ≥ 0.07. **D,** Genus-level fungal community abundances in winery samples, threshold RA ≥ 0.055.

### Indoor environmental conditions are associated with microbial community composition

Next, we sought to determine whether environmental conditions within fermented food production facilities could be linked to spatial and temporal changes in microbial communities on equipment and environmental surfaces. Remote sensors were placed in different production areas in each of the five food facilities, as described in a previous study (40). These sensors continuously monitored atmospheric temperature, relative humidity (RH), and carbon dioxide (CO_2_) concentration at each site, and volatile organic compound (VOC) concentration at sites located in the primary fermentation areas (Figure S2). As described previously, fluctuations in indoor atmospheric conditions reflect a combination of seasonal changes, indoor climate control (in the case of some facilities), and food production practices (40). Outdoor spaces and barrel rooms, the latter which typically have a high-RH controlled atmosphere, had higher RH than indoor spaces. In wineries, CO_2_ and VOC concentrations spiked around harvest periods, when active wine fermentations were emitting these compounds into the indoor atmosphere (Figure S2). These patterns are not observed in Winery 1 (Figure S2) because each fermenter is equipped to emit headspace vapors into a manifold that vents to the outside of the facility, reducing the need for indoor ventilation (40). Similarly, fluctuations in CO_2_, VOC, and RH in the creamery aging rooms (Figure S2) mirrored the progression of cheese fermentations in these facilities (40).

Hence, in both creameries and wineries, the main fermentation areas tended to have higher means and variances for CO_2_ and RH than in non-production areas (e.g., laboratories and raw material handling areas). Given these temporal characteristics, we chose six environmental measurements to correlate with microbial measurements: daily means and weekly coefficients of variation (CV) for temperature, RH, and CO_2_. Since VOC was only measured at select sites, it was excluded from microbial association tests.

We performed PERMANOVA tests and PCoA to assess whether indoor environmental measurements were associated with microbial beta diversity (Tables 1-4, Figure 3). Significant results with these tests indicate that indoor conditions correlate with bulk changes in microbial populations, and hence that large-scale patterns of microbial community composition may be influenced by indoor environments. Overall, environmental measurements were frequently found to be significantly associated with beta diversity (P < 0.05), though the contribution of indoor environment to sample variation was much less than those of facility and sample type (net R^2^ ≤ 0.09) (Tables 1-4). In general, creamery microbiota displayed fewer significant associations with environmental conditions (Tables 1 and 3; N = 6 factors with P < 0.05) than winery microbiota (Tables 2 and 4; N = 27 factors with P < 0.05), perhaps because stricter climate control within these facilities diminished potential variation.

**Figure 3.**
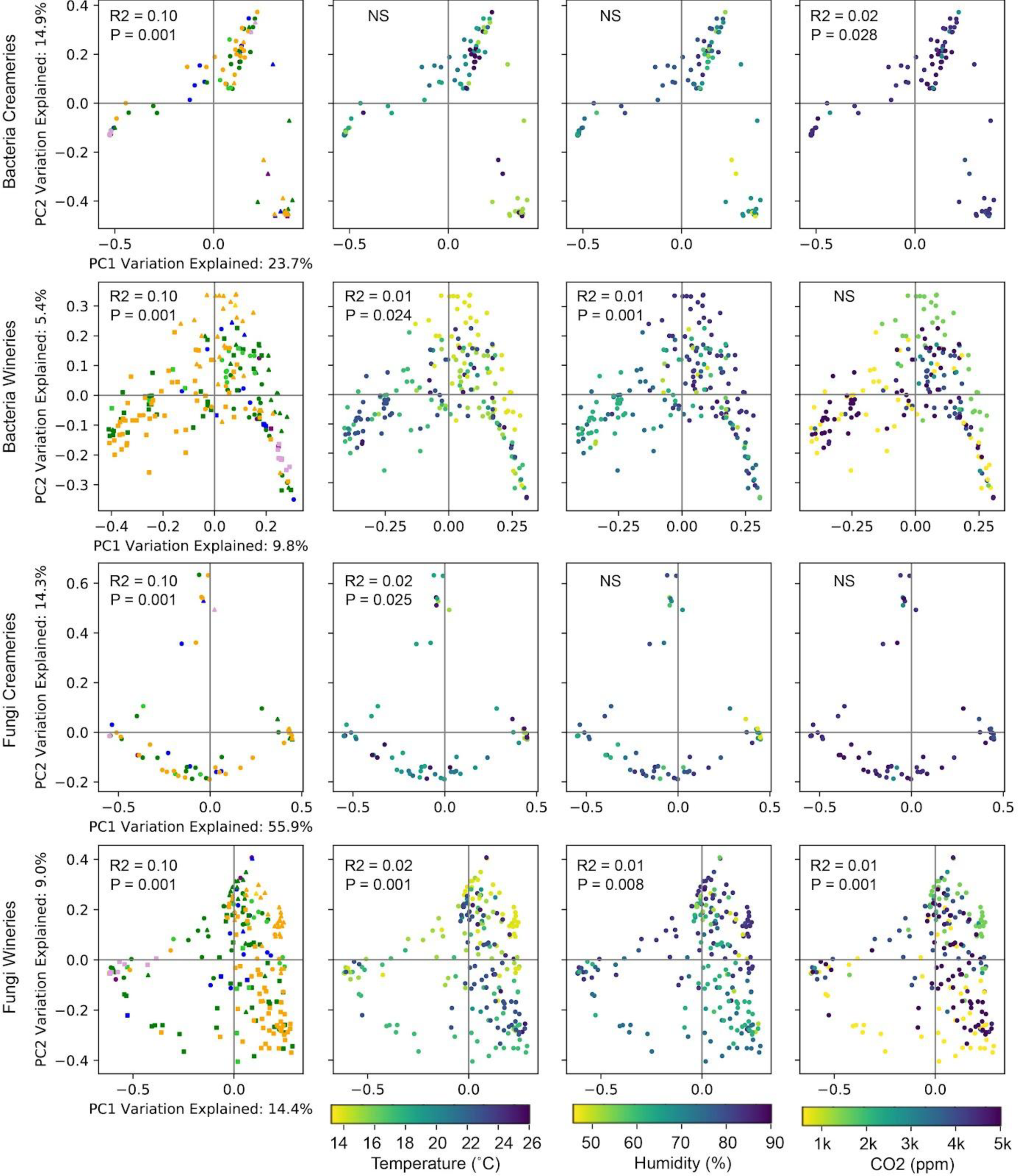
Bray-Curtis dissimilarity principal coordinate analysis visualizes relationship between indoor environmental conditions and microbiome. PERMANOVA R^2^ and P-values from Tables 1-4 are displayed for each panel (if P < 0.05). NS = not significant (P > 0.05). See Figure 1 for a sample type color key for the left-most column of subplots. All other columns are color-coded by the magnitude of the environmental measurement (Temperature, Relative Humidity, CO2) noted at the bottom of all graphs in a single column.

In creameries, CO_2_ displayed the strongest associations: daily mean CO_2_ was significantly associated with bacterial Bray-Curtis dissimilarity (R^2^ = 0.02, P = 0.028) and Jaccard distance (R^2^ = 0.02, P = 0.002), and weekly CO_2_ CV was significantly associated with bacterial Bray-Curtis dissimilarity (R^2^ = 0.02, P = 0.018) and weighted UniFrac distance (R^2^ = 0.02, P = 0.015) (Table 1). Creamery daily mean RH was also significantly associated with bacterial Jaccard distance (R^2^ = 0.02, P = 0.010) (Table 1) and daily mean temperature with fungal Bray-Curtis dissimilarity (R^2^ = 0.02, P = 0.025) (the only measurement associated with fungal beta diversity; Table 3).

Wineries displayed much stronger associations between microbial beta diversity and indoor environmental conditions. Almost all environmental measurements were significantly associated with both bacterial and fungal beta diversity (P < 0.05), with the exception of weekly CV temperature, which was only significantly associated with fungal Bray-Curtis dissimilarity (R^2^ = 0.01, P = 0.036) (Tables 2 and 4). These differences are apparent in PCoA scatterplots, in which gradients can be observed between high and low values of mean temperature, RH, and CO_2_ (Figure 3), indicating that environmental conditions are associated with large-scale differences in bacterial and fungal communities at these sites. Hence, it appears likely (if still correlative) that local changes in indoor environment impact microbial communities in food production facilities; climate control (as practiced throughout the creameries) reduces this impact but is nevertheless expected to impact community structure.

Alpha diversity (within-sample diversity) is another useful measurement of microbial diversity within communities. We measured the observed number of unique bacterial and fungal amplicon sequence variants (ASVs) in each sample, as an indicator of microbial richness. Analysis of variance (ANOVA) tests were performed to determine whether facility, sample type, room type, and environmental factors impact species richness. Facility and room type did not exhibit significant effects, but sample type significantly impacted bacterial and fungal richness in both creameries and wineries, with the exception of creamery fungi (P < 0.05, Figure S3, Table S6). Higher daily mean temperatures and higher daily mean RH were both separately associated with significantly lower bacterial diversity in both creameries and wineries (Table S6).

### Indoor environmental conditions are predictive of species abundances in production environments

Environmental conditions appear correlated with large-scale differences in microbial compositions in both wineries and, to a lesser extent, creameries. As a next step, we used multiple methods to explore associations between environmental conditions and the relative abundance of individual organisms at different sites. These all represent correlative, exploratory analyses; they do not imply causation and hence we did not attempt to determine probability values. Instead, these analyses should be considered hypothesis-generating, and will allow follow-up experiments to verify causative and mechanistic relationships.

Canonical correspondence analysis (CCA) is a multivariate constrained ordination method that tests for correspondence between a matrix of explanatory (environmental) and response (microbial) variables (41). CCA biplots (with type II scaling) identify several associations between environmental conditions and facility microbiota (Figure 4). Clear bacterial community trends can be observed in creameries: CO_2_ daily mean, CO_2_ weekly CV, and RH daily mean all correlate positively with a group of ASVs identified as belonging to class *Clostridia*, and negatively with diverse *Alphaproteobacteria* and *Gammaproteobacteria* (Figure 4A). A cluster of *Pseudomonas*, *Enterococcus, Lactococcus*, and *Vibrio* ASVs positively corresponds with daily mean temperature (Figure 4A center-right), RH, and CO_2_, while a separate cluster of *Psychrobacter, Pseudomonas*, *Bacillus*, and *Lactobacillus* ASVs negatively correspond with daily mean RH and CO_2_, positively with weekly CV of temperature and RH, and at low to moderate values of daily mean temperature (Figure 4A, lower-left corner and left edge). A large number of *Brevibacterium* and *Brachybacterium* ASVs, found abundantly on some types of aging cheeses in these creameries (5), are associated with low daily mean temperatures but vary along a gradient of daily mean RH and CO_2_ values (Figure 4A left edge). Creamery fungi do not display such a clear taxonomic distribution, though a range of yeasts are found associated with low daily mean temperatures, RH, and CO_2_, but high weekly RH CV (Figure 4B).

**Figure 4.**
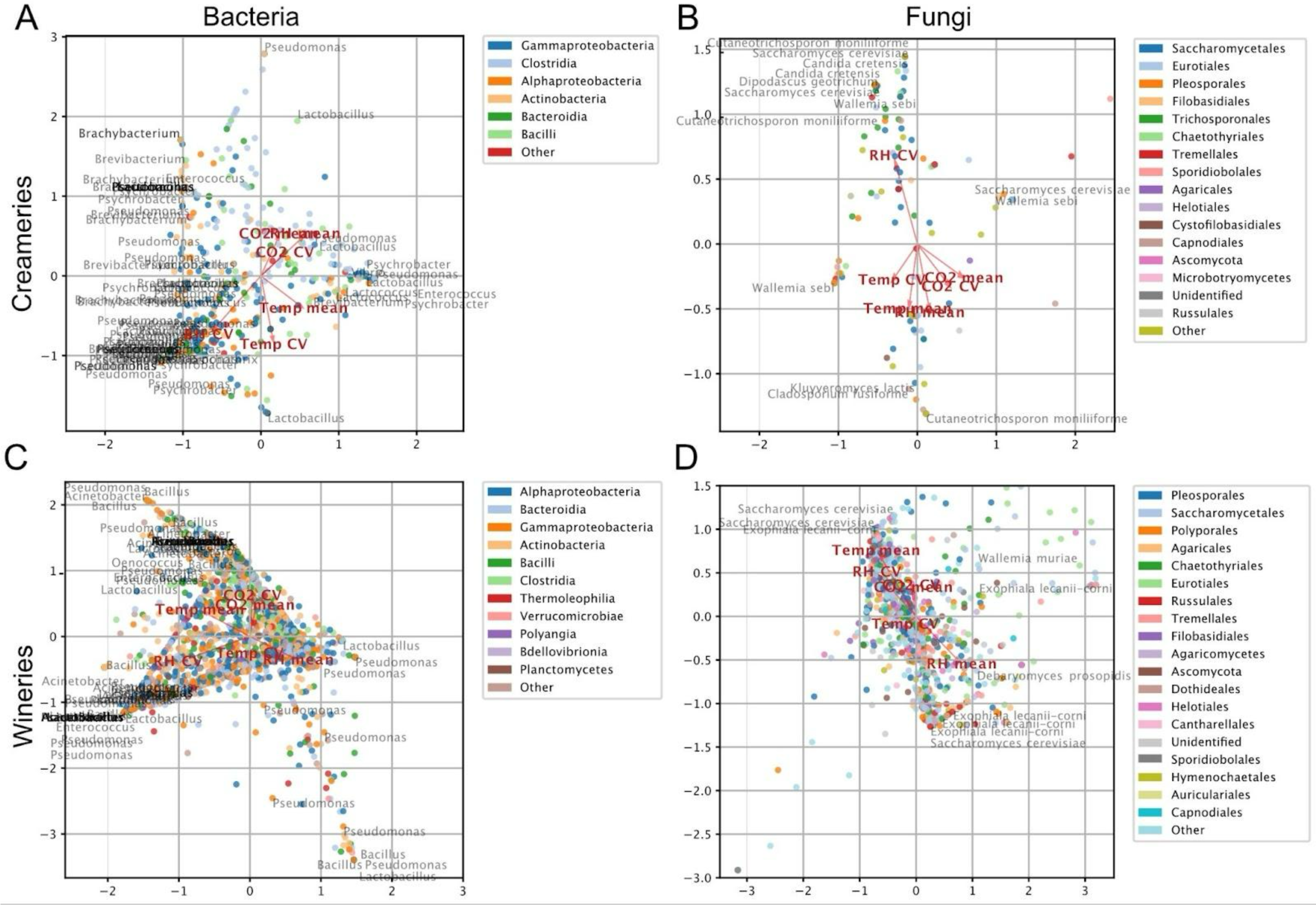
Canonical correspondence analysis ordination biplots detect associations between indoor environment and microbial community composition. Canonical correspondence analysis of daily mean temperature (Temp mean), humidity (RH mean), and CO_2_ (CO_2_ mean) and weekly coefficient of variance for temperature (Temp CV), humidity (RH CV), and CO_2_ (CO_2_ CV) as explanatory variables; and bacterial ASV (left panels) or fungal ASV community compositions (right panels) in creameries (top panels) or wineries (bottom panels) as observations. Each panel displays the co-ordination of variables along CCA axes 1 (horizontal) and 2 (vertical). Only select ASVs with strong associations are labeled (via species label) in each panel (genus/species labels are placed at approximate positions to increase readability). Points (ASVs) in each plot are color-categorized by class (for bacteria) or order (for fungi), or the next-lowest rank if that ASV is incompletely classified. Note that panels B and D are truncated to exclude a small number of outliers.

Winery bacteria and fungi display widely variable associations within taxonomic clades (Figure 4C, D). Various ASVs of the bacteria *Lactobacillus*, *Bacillus*, and *Pseudomonas* and the fungi *Saccharomyces cerevisiae*, *Exophiala lecanii-corni*, and *Wallemia muriae* are associated more strongly with daily mean temperature, RH, or CO_2_ (Figure 4C, D). Among these, *Lactobacillus* and *S. cerevisiae* are prominent fermentative organisms for malolactic and alcoholic fermentation, respectively, but whether these ASVs relate to strains dominant in the wine fermentations conducted at these wineries is not known; nevertheless, these findings indicate that indoor conditions may influence the selective establishment and retention of different fermentative strains within wineries, indirectly impacting wine quality and contamination risks.

Multinomial regression reference frames, a statistical method for determining differential abundance from relative abundance data (42), provides an alternative approach for estimating associations between explanatory (environmental) and response (microbial) variables. Multinomial regression biplots (Figure S4) illustrate taxonomic associations with environmental conditions, and differential rank heatmaps identify the ASVs most strongly associated with each environmental variable (Figure 5). In the creameries several *Vibrio, Pseudoalteromonas*, and *Halomonas* ASVs – all halophilic bacteria associated with daily mean temperature, RH, and CO_2_ (Figure S4A left side) – are among the most strongly associated with these variables (Figure 5A). A small number of creamery fungi display significant associations (Figures 6B and S3B), consistent with the very weak effects of environmental conditions on creamery fungal beta diversity (Table 3). However, these include the important cheese-fermentative fungi *Dipodascus* and *Debaryomyces* species (Figures 6B and S3B). *Dipodascus geotrichum* (teleomorph of *Geotrichum candidum*) is positively associated with temperature and RH mean and CV, whereas two different ASVs classified as *Debaryomyces prosopidis* demonstrate opposing associations with these variables (Figure 5B). In wineries, multinomial regression mirrors the results of CCA, with no particular pattern to taxonomic associations (Figure S4C-D); the top differential ranks among bacteria are primarily non-fermentative (Figure 5C), but several fermentative fungi are among the top differential ranks: *S. cerevisiae* ASVs associated positively with mean daily temperature, and *Saccharomyces kudriavzevii* and *Pichia* ASVs negatively associated with mean daily temperature and RH (Figure 5D). Notably, *S. kudriavzevii* is commonly described as a cryotolerant yeast which grows better at lower temperatures, (43, 44) which corroborates the association seen here.

**Figure 5.**
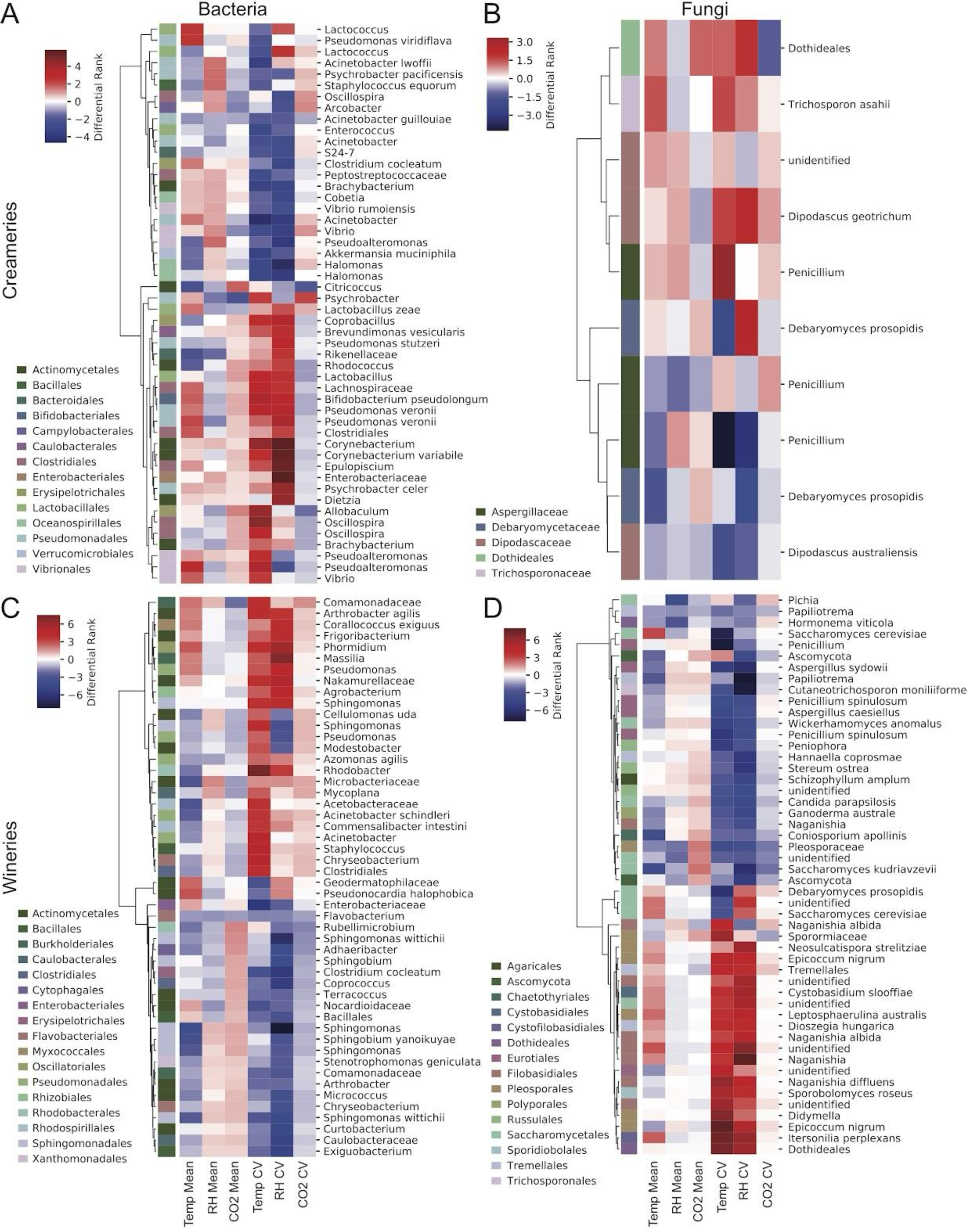
Multinomial regression differentials heatmaps indicate associations between indoor environment and microbial community composition. Multinomial regression of daily mean temperature (Temp mean), humidity (RH mean), and CO_2_ (CO_2_ mean) and weekly coefficient of variance for temperature (Temp CV), humidity (RH CV), and CO_2_ (CO_2_ CV) as explanatory variables; and bacterial (left panels) or fungal community compositions (right panels) in creameries (top panels) or wineries (bottom panels) as observations. Each panel displays cluster maps of the top 50 highest and lowest ASVs (ranked by relative differential score) for each dataset; cladograms represent UPGMA hierarchical clustering of each ASV as a function of their differential rank score similarities, measured by pairwise Bray-Curtis dissimilarity. The differential rank indicates the strength of association between each taxon and environmental variable. ASVs (rows) are labeled by species name or the next-lowest taxonomic rank if that ASV is incompletely classified. Row margins are color-coded by taxonomic order, as indicated by the color key.

We used Random Forest regressors (45) in q2-sample-classifier (46) as a third method to explore associations between environmental conditions and microbial communities. This approach trains a machine-learning regressor to predict a continuous variable (in this case, the daily mean or weekly CV of some environmental factor) and identify which microbial features are most predictive. If the microbiome is predictive of an environmental condition it suggests that the environment may be associated with the temporal composition of the microbiome at that site. Each regressor was trained to predict a single variable in either wineries or creameries, and each was only weakly or moderately predictive of environmental characteristics (Table S7). Mean daily temperature and weekly CV humidity models exhibited the best predictions in both wineries and creameries, as judged by low mean squared error (MSE) values and moderate R^2^ and slope values for correlation between predicted and true values (Table S7). In creameries, the most predictive features (i.e., the microbes that are most strongly associated with the target variable) include a number of important fermentative and spoilage organisms, including *Psychrobacter, Lactococcus, Dipodascus geotrichum,* and *Debaryomyces* yeasts (Figure S5). In wineries, the most predictive features are primarily non-fermentative organisms with no known links to wine quality (Figure S5). The predictive strength of these single-target models is likely weakened by the multiple variables at play that impact microbial community structure on different surface types throughout the facilities, as evinced by the high residual values in the multivariate PERMANOVA tests (Tables 1-4). Nevertheless, the top important features identified by these models highlight potential connections between indoor climate and microbial community assembly, with potential impacts on food quality characteristics.

### Survey of spoilage organisms and potential foodborne pathogens in dairy facilities

Given the sensitivity of dairy processing facilities to microbial spoilage and safety risks, we performed a survey of potential foodborne pathogens and dairy spoilage bacteria within the dairy facilities. It is critical to understand that the methods we employed here cannot detect strain-level differences or infer the presence of pathogenesis/virulence-related genes necessary to positively identify a pathogen. Many of these genera contain non-pathogenic species and strains, and others (e.g., *Staphylococcus*) also contain species that are considered beneficial non-starter strains in production of some cheeses; as such, this approach should be seen as purely exploratory. Moreover, since these methods do not differentiate viable from non-viable organisms, it is possible that DNA found in the environment may represent DNA from dead cells, e.g., of raw milk bacteria that are killed by pasteurization.

We filtered results for the presence of 16 genera that include potential foodborne pathogens, focusing on those with relevance to dairy fermentations: *Listeria, Bacillus, Yersinia, Salmonella, Escherichia, Campylobacter, Clostridium, Staphylococcus, Mycobacterium, Francisella, Brucella, Aeromonas, Coxiella, Shigella,* and *Vibrio*. None of the principal dairy-borne pathogenic genera – being *Listeria, Yersinia,* and *Salmonella* – were detected in this study. With the exception of *Vibrio* and *Staphylococcus*, ASVs assigned to other potentially pathogenic groups were detected at very low mean frequencies (Table 5). None of these ASVs (again, with the exception of *Vibrio*) displayed any significant associations with indoor environmental conditions (Figure 5A).

**Table 5.** Distribution characteristics of potential human pathogenic genera and species detected in creameries.

| Genus | Mean (%) | Median (%) | Max (%) | ASVs | Samples |
| --- | --- | --- | --- | --- | --- |
| *Mycobacterium* | 0.001 | 0.000 | 0.297 | 18 | 39 |
| *Campylobacter* | 0.000 | 0.000 | 0.023 | 15 | 15 |
| *Bacillus* | 0.001 | 0.000 | 0.220 | 37 | 60 |
| *Staphylococcus* | 0.025 | 0.002 | 1.000 | 30 | 302 |
| *Clostridium sensu stricto 1* | 0.001 | 0.000 | 0.025 | 19 | 57 |
| *Clostridium sensu stricto 12* | 0.000 | 0.000 | 0.001 | 1 | 1 |
| *Clostridium sensu stricto 13* | 0.000 | 0.000 | 0.075 | 7 | 27 |
| *Clostridium sensu stricto 7* | 0.000 | 0.000 | 0.001 | 1 | 3 |
| *Clostridium sensu stricto 8* | 0.000 | 0.000 | 0.000 | 1 | 1 |
| *Aeromonas* | 0.001 | 0.000 | 0.019 | 3 | 99 |
| *Coxiella* | 0.000 | 0.000 | 0.004 | 3 | 6 |
| *Escherichia-Shigella* | 0.002 | 0.000 | 0.074 | 10 | 71 |
| *Francisella* | 0.000 | 0.000 | 0.001 | 1 | 1 |
| *Vibrio* | 0.003 | 0.000 | 0.152 | 14 | 123 |

\* ASVs = number of ASVs classified to that genus/species. Samples = number of samples in which that genus/species was observed. Mean, Median, and Maximum represent relative abundances as a percentage. “Unclassified” species could not be confidently classified at species level; “unknown” species received a species-level classification to a reference sequence group without a species-level annotation.

The potential pathogen groups that were detected (Table 5) were only present at low prevalence and low frequency on non-processing surfaces (e.g., drains, floors). *Campylobacter* ASVs were detected at low frequency in two rennet samples (< 0.5% relative frequency), an unwashed cheese mold (0.08% relative frequency), and on restroom surfaces (as high as 2% relative frequency on a sink handle). *Clostridium* ASVs were detected in three rennet samples (as high as 0.5% relative frequency) and two young cheese samples (0.1% relative frequency in both). *Mycobacterium* ASVs were detected in floor drains and sinks, consistent with frequent reports of *Mycobacterium* spp. in water distribution systems (47). *Shigella* spp. were detected in a single cheese shelf swab (0.2% relative frequency) and on a restroom soap dispenser (1.6% relative frequency). *Coxiella* ASVs were found in four raw milk samples at low frequency (< 0.1% relative frequency). A *Francisella* sp. was found in a single river water sample collected near one facility, but it was not detected within any of the facilities sampled. It is important to highlight that short, V4-domain 16S rRNA gene sequences are not sufficient for definitively identifying species (classification with SILVA is only performed here to the genus level), nor for predicting the pathogenicity of a given species, and that detection from DNA (as performed here) does not differentiate live vs. dead biomass. Hence, this exploratory analysis cannot be used to confirm the presence of human pathogens in the facilities, or predict any foodborne hazards.

The *Vibrio* sequences detected here most likely do not belong to pathogenic species; of the 20 ASVs classified as *Vibrio*, a BLAST search against NCBI RefSeqs yielded top hits to *Vibrio casei* (among other species with ≥98% similarity), which has recently been isolated from various cheeses (48). Similar *Vibrio* ASVs have also been detected on cheeses in several molecular and marker-gene-sequencing-based studies (49–51). Hence, the *Vibrio* sequences detected here are most likely benign, and may actually contribute to the sensory characteristics of the cheeses, given their relatively high abundance (Table 5). We have previously identified similar *Vibrio* ASVs to be a component of the “house microbiome” in cheese facilities (5, 51). These and the other halophiles detected on cheeses and in the environment in this study, such as *Halomonas* and *Pseudoalteromonas*, may be introduced during brining and might contribute sensorily to these cheeses (51).

Next, we screened the data for potential spoilage bacteria in the cheesemaking facilities, focusing on the following genera: *Psychrobacter, Pseudomonas, Brochothrix, Alcaligenes, Enterobacter, Citrobacter,* and *Serratia* (Table 6). Several species of *Psychrobacter, Pseudomonas,* and *Brochothrix* were widespread within the facilities, though typically at very low frequencies. *Pseudomonas* was the most abundant among these, detected in 368 samples at as much as 98.8% relative frequency (in a raw cream sample); this taxon was also detected abundantly in a raw milk sample (77.2%) and on uncleaned equipment (e.g., 26.9% relative frequency on a draining table; 36.4% in a condensation trap; 33.2% on a squeegee), but it was most frequently found on floors and drains. *Psychrobacter* species were detected frequently on floors, used processing equipment, and several cheeses, including one spoiled cheese sample on which *Psychrobacter* (unknown species) was detected at 91.3% relative abundance. *Brocothrix* was detected in raw cream (at up to 5.5% relative abundance), raw milk (max 1.8% relative abundance), used processing equipment, and cheese samples (max 0.3% relative abundance). *Alcaligenes* was detected at low abundance (mean 0.02% relative frequency) and prevalence (7 samples), primarily on floors in the draining room. *Serratia* spp. were detected at low abundance (mean 0.01% relative frequency) and prevalence (15 samples) in drains and floor samples, as well as in a rennet paste sample (0.1% relative frequency). *Enterobacter* spp. and *Citrobacter* spp. were not detected in any samples.

**Table 6.**
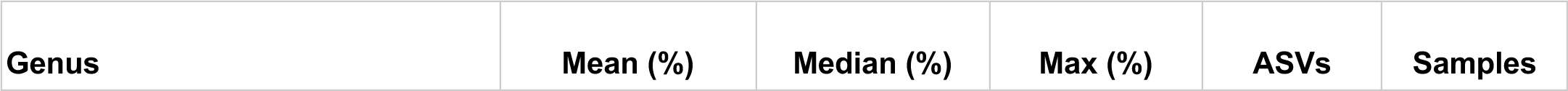
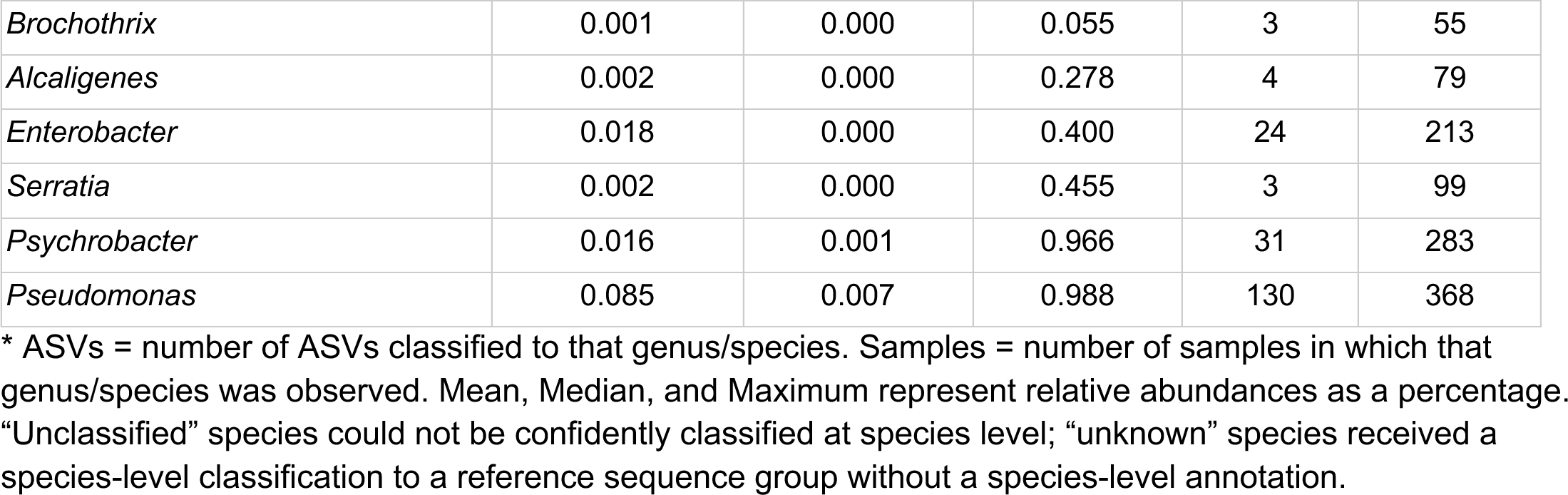
Distribution characteristics of potential dairy spoilage bacteria detected in creameries.

## Discussion

Food production environments can serve as a reservoir for food spoilage organisms and potential human pathogens (52), but also for non-starter organisms that beneficially contribute to the quality characteristics of many fermented foods (16, 53, 54). The challenge facing food producers who rely on adventitious microbial activity in their foods is to maximize the beneficial contributions of some non-starter organisms while simultaneously shielding foods from exogenous food-spoilers and pathogens via routine cleaning, sanitation, and hygiene surveillance. Indoor climate control offers one mechanism by which food producers can manipulate microbial risks within food production facilities, as is already commonly practiced through temperature and condensation control. The broader impacts of indoor environmental conditions on microbial community compositions within food production facilities have been little explored.

This study demonstrates that indoor environmental conditions are associated with the spatial distribution and temporal fluctuations of microbiota within food production facilities, suggesting that the indoor environment is directly influencing these microbial communities. We tested our hypothesis within five fermented food production facilities, comprising two different types of food producers (wineries and cheese makers) to confirm that our observations generalize across multiple sites and production conditions. We draw several conclusions from our findings. First, facility and sample type exert stronger effects on microbial composition than any individual indoor environmental factor – though combined environmental effects exert a similar degree of influence on microbial diversity (Tables 1-4). This is not a surprising result, as we and others have shown previously that different creameries possess a unique “house microbiota” of non-starter microbes which are abundant on the cheese surface and distinct to individual facilities (5, 55) – though there is also evidence that such between-facility microbial differences may be due at least in part to daily within-facility microbial variations (56). We expect that the inter-facility effect observed here is common to a variety of food production facilities, given that regional factors can strongly influence the microbiota of raw materials (56–58), altering the microbial inputs to food facilities and directing the course of microbial succession during food fermentation (11, 54–56, 59). This study represents a foundational attempt at quantifying the effects of facility-specific differences in indoor environment on microbial inhabitants in fermented food production facilities, with implications for facility design and indoor climate control.

We found that several aspects of indoor environment are associated with spatial and temporal differences in microbial communities and specific microbial taxa. In some cases, this reflected seasonal changes in ambient conditions, making it difficult at times to untangle natural from anthropogenic effects. For instance, in wineries, changes in facility environmental conditions were correlated with different stages of the harvest and production schedule. In other cases, fluctuations in indoor conditions were clearly related to production schedule and human activity. For example, weekly oscillations in humidity occurring during dry seasons in California can be linked to increased water use within the wineries on weekdays (e.g., for cleaning equipment and winery surfaces) as compared to on weekends, when work is usually paused or reduced (40).

Both natural and anthropogenic effects on the indoor environment can be mitigated through climate control, and potentially through changes in standard operating procedures. For example, we identify several species of the potential dairy-spoiling bacterium *Psychrobacter* associated with high RH in creameries (Figure 5A); humidity control and/or changes to washing protocols may provide an avenue for diminishing *Psychrobacter* contamination in dairy facilities. However, this requires additional investigation to validate. Interestingly, we detected several taxa that were associated with rapidly fluctuating environmental conditions, including spoilage bacteria such as *Psychrobacter* spp. and *Pseudomonas* spp., which in creameries were associated with high weekly CV in temperature and RH (Figure 5A). More research is required to determine the mechanism and biological significance behind these associations, but it is possible that these organisms are able to flourish under highly variable conditions, under which other organisms are inhibited by temperature and humidity extremes. Indeed, *Psychrobacter* is psychrotrophic and thought to be somewhat halophilic,(53, 60) which could explain in part its tolerance to changes in temperature, humidity, and other environmental conditions. The species of this genus, along with *Pseudomonas* species, are found nearly ubiquitously in both food facilities and natural environments; as such, members of these genera may be particularly able to adapt to a variety of environmental conditions.(61)

Our exploratory analysis identified several novel associations between indoor environmental conditions and quality-related organisms, which require further investigation to validate. In some cases, there was a clear trend between taxonomic affiliation and environmental niches: for example, in creameries, *Brevibacterium* and *Brachybacterium* ASVs were associated with relatively low to moderate temperatures and RH, and *Lactococcus* ASVs were associated with relatively high temperatures and RH (Figure 4A). Both *Brevibacterium* and *Brachybacterium* species are routinely found on cheese rinds and ripening boards,(54, 55, 59, 60, 62–64) developing over long periods of time in low-temperature ripening rooms. Furthermore, the application of saline solutions to the cheese rind reduces water activity, selecting for tolerant species.(62) As such, the association of these genera with low temperatures and RH align with the environments in which they tend to be found. However, ripening rooms themselves do tend to be humid, and *Brevibacterium* has been reporting as only growing at RH >90% (65) with some species growing better at higher levels of RH and temperature,(66) so the associations seen in this work may not be definitive for the entire genus and/or merely reflect the relative tolerance of these genera. Furthermore, both *Brevibacterium* and *Brachybacterium* have been found in water samples from dairy farms and cheese facilities, with a notable dominance of the former genus (67), which suggests their ability to survive, if not necessarily thrive, in high-water-activity environments. As for *Lactococcus*, it is a common starter bacteria used in cheese making.(56, 59) It has been found to dominate microbial communities at the bottoms of wooden vats used for ripening milk overnight, as well as dominating ripening and finished cheese interiors.(53, 54, 56) Ripening vats can be quite humid environments, at least when they are in use, which could explain the association between *Lactococcus* and high RH in this study. *Lactococcus* species have also been found in the water of both dairy farms and dairy facilities.(67) The *Lactococcus* species used in cheesemaking are known to grow optimally around 30°C,(68) which aligns with its associations with higher temperatures in this study. However, since *Lactococcus* is generally prevalent in creameries and creamery/dairy products,(53, 55, 56, 67) it may be capable of growing in myriad environmental conditions.

In many other cases, different variants/subspecies (ASVs) of the same quality-related taxa displayed opposing associations with environmental conditions. For example, some ASVs of the psychrotolerant dairy-spoilage bacterium *Psychrobacter* were strongly associated in creameries with relatively high temperatures, and others with relatively low temperatures (Figures 5A and 6A). It should be noted, however, that this genus was also associated with cooler temperatures on wooden cheese boards in another study.(62) *Pseudomonas* ASVs in creameries formed three groups: those associated with relatively high temperature and RH, those associated with relatively moderate to high temperature and low RH, and those associated with relatively low temperature and RH (Figures 5A and 6A). As previously mentioned, the diversity of environments in which *Psychrobacter* and *Pseudomonas* are found suggest their resilience to environmental changes and potential intra-genus phenotypic diversity.(61) Similar reasoning could explain the opposing associations in this work between various *Debaryomyces* ASVs and daily temperatures and RH – this genus is commonly found in myriad low moisture environments, including cheese, and has been described as an extremophile and a “metabolically versatile” genus with intra-species phenotypic diversity.(69)

In wineries, two ASVs of the wine yeast *Saccharomyces cerevisiae* were associated with relatively high temperatures and low RH (Figures 4D and 5D), while a third was associated with relatively low temperatures and high RH (Figure 4D). Though *S. cerevisiae* is usually described as growing better at higher temperatures – at least compared to non-*Saccharomyces* or non-*cerevisiae* yeasts – (43, 70, 71) there have also been reports of *S. cerevisiae* strains displaying wide ranges of temperature preferences, or with tolerance to a broad range of growth temperatures (43, 71). This could help explain the varied associations seen here between *S. cerevisiae* ASVs and daily mean temperature. More work needs to be done to determine how these opposing distribution patterns may relate to microbial phenotype and spoilage risks. For example, different strains of *S. cerevisiae* can serve as beneficial fermentative organisms or as spoilage yeasts in beverage fermentations (3); if their strain phenotypes (e.g., preferring higher vs lower temperatures) correlate with their environmental niche, the impact of indoor environment on their ecological distribution has important implications for quality control.

Similarly, it is unclear whether the potential spoilage bacteria that were widespread in the creameries (e.g., *Psychrobacter* spp. and *Pseudomonas* spp.) actually pose a risk: some members of these genera are dairy spoilers, but most species are widely distributed in the environment and are not known to cause food spoilage (61, 72, 73). It will be essential to study the spoilage potential of individual strains in order to evaluate whether their ecological niche correlates with their spoilage phenotype.

This leaves several open questions that are ripe for future investigation, as well as some limitations of the current work. For instance, the associations discovered in this study are correlative, and no causation is implied. Furthermore, the sequencing methodology used here measures relative abundances of taxa, and does not differentiate between live and dead cells; as such, it is unclear whether the observed changes in taxon abundances relate to actual changes in their biomass or activity. Future work will build on these findings to test whether manipulating environmental conditions can support desired changes in microbial community structure and activity. An additional direction for future work will be to examine how ecological interactions among microbial community members may impact the retention of beneficial or detrimental populations in the studied environments and environmental conditions. Another might be to further quantify the extent and duration of surface wetness, since these will be related to evaporation and drying rates and wetness due to condensation and washdown frequency. Sensors could be installed to follow these conditions but they would need to be more robust than those used in this study. Future work will examine the interaction networks present within food production facilities, especially how these networks may be leveraged to suppress the establishment of pathogens and food-spoilage microbes on food surfaces.

Overall, these findings have clear implications for the design and operations of these facilities in terms of the environmental temperature and humidity in each processing room or space. While this study used the primary air properties of bulk air temperature, humidity, and CO2 levels, the important variables for designers in the future may instead include the dew point of the air in the room and the surface temperatures that would establish the incidence of condensation and the time of wetness on these surfaces. These conditions also interface with washdown and cleaning practices, surface areas in proportion to the volume of the space, and the intake of outside air and its condition. The bulk conditions are expected to be more regular throughout the year in the creameries than in wineries, due to climate control measures routinely used in the former, and seasonal production schedules in the latter. The role of outside air and its conditioning will vary between facilities, their locations and the season.

In conclusion, we show for the first time the dynamic relationship between indoor environmental conditions and the composition of bacterial and fungal communities across space and time in food facilities. We show that the primary microorganisms present on processing and non-processing surfaces are largely related to the microbiota of the raw materials and fermented products, and that their community assembly shifts over time and space, significantly associated with temperature and humidity gradients within each facility. Although significant, individual conditions were only weakly to moderately predictive of microbial community composition at a given sampling site, suggesting that complex interacting factors (including conditions not monitored in this work) shape microbial presence on food processing facility surfaces. More work is needed to enable better predictive control of microbial communities within food processing facilities. Improved understanding of how the built environment shapes microbial community dynamics within food facilities will enable rational design of facilities in the future that passively control microbial growth for improved food quality and safety.

## Methods

### Sample Collection

Samples were collected from two commercial cheese makers (creameries) and three wineries in the Northern California area. Surfaces were sampled using sterile cotton swabs dipped in sterile-filtered 1X PBS (phosphate-buffered saline) to swab a ∼25.8 cm^2^ (4 in^2^) surface area as described previously (12). Samples were stored in sterile tubes, and immediately transported to the lab for storage at -80C prior to analysis. Other samples, including liquid or solid materials, e.g., wine, water, grape must, were aseptically sampled and directly placed in sterile containers. Air samples were collected using 20mL of sterile-filtered 1X PBS in a Biosampler (Cat.# 225-9594 and #225-9596, SKC Ltd.,PA, USA).

### Amplicon library preparation and sequencing

DNA extraction, amplification and sequencing was performed using Earth Microbiome Project (EMP) protocols as previously described (https://earthmicrobiome.org/protocols-and-standards/) (74). Briefly, DNA was extracted from environmental swab samples with the MoBio PowerSoil-htp 96-Well DNA Isolation Kit (12955-4, MoBio Laboratories, Inc., Carlsbad, CA, USA) according to the manufacturer’s instructions. Sample homogenization was carried out on a Geno/Grinder 2010 tissue homogenizer. Amplification was carried out in triplicate of the V4 16S rRNA gene with 515f (Parada)/806R(Apprill) primers (75, 76) and of the ITS region of fungal DNA with ITS1f-ITS2 primers (77), both sets contained 12bp Golay error-correcting barcodes on the reverse primers and were constructed as described in the EMP protocols. Amplicons were run on an 0.8 % agarose gel to verify amplification by gel electrophoresis. Bacterial and fungal amplicons were combined into two separately pooled samples, purified using the Qiaquick spin kit (Qiagen), and submitted to the University of California Davis Genome Center DNA Technologies Core for Illumina 250-bp paired-end sequencing on an Illumina MiSeq instrument.

A total of 2,329 samples were profiled using 16S rRNA gene amplicon sequencing, in addition to 124 negative controls included on each plate (swab blanks and DNA sequencing kit and PCR negative controls) (2,453 total samples including blanks). A total of 2,446 samples were profiled with fungal ITS domain amplicon sequencing, including 119 negative controls (swab blanks and DNA sequencing kit and PCR negative controls).

### Bioinformatics

Sequence data were processed and analyzed using the microbiome bioinformatics framework QIIME 2 (78). DADA2 (79) was used (via the q2-dada2 QIIME 2 plugin) to filter and correct erroneous sequences, remove PhiX and chimeric reads, and merge paired-end reads. The ITS sequences were too short to overlap paired-end reads after denoising, so only forward reads were used for downstream analysis. The plugin q2-cutadapt (80) was used to trim primers and adapters from the ITS sequence data prior to denoising. Taxonomy was assigned to sequence variants using q2-feature-classifier (81) with the classify-sklearn method using a pre-trained Naive Bayes classifier trained for 16S rRNA gene sequences against Silva release 138.2 (82) with reference sequences clustered at 99% similarity and weighted by average taxonomic frequencies (83); against the UNITE version 7.1 release (84) (for fungal ITS sequences); or Greengenes v.13_8 (85) (for initial classification and filtering of chloroplast and mitochondrial sequences from 16S rRNA gene data). To construct a phylogenetic tree, denoised 16S rRNA gene amplicon sequence variants (ASVs) were grafted into the SILVA reference tree with the QIIME 2 plugin q2-fragment-insertion (86). K-mer frequencies were calculated with feature tables filtered to 1,000 sequences per sample using q2-kmerizer in QIIME 2 with default settings and a maximum of 5,000 features (38). These feature tables either contained all sample types except for outdoor and building air (for global PERMANOVA analyses), or only contained the following sample types (for pairwise PERMANOVA analyses): raw materials, fermented food products, processing equipment, and building surfaces.

Built environment samples are relatively low-biomass, increasing the risk of exogenous contamination and cross-contamination. We employed multiple types of control samples (swab, DNA extraction kit, and PCR blanks) and multiple filtering methods to remove potential contaminants. Feature tables were filtered using the R package decontam (87) to remove putative reagent contamination and exogenous contaminants detected in the negative controls. The QIIME 2 plugin q2-feature-table was used to filter other non-target reads, including sequences taxonomically classified as mitochondrial or plastid DNA, and sequences that did not receive at least phylum-level classification. After filtering to remove contaminants and samples with insufficient read depth (<1,000 sequences per sample), 1,694 samples (including 8 blanks) remained in the bacterial 16S rRNA gene dataset, and 1,694 (including 6 blanks) in the fungal ITS dataset. Thus, 27.6% of biological samples and 93.5% of blanks were filtered from the bacterial 16S rRNA gene dataset, and 27.5% of biological samples and 95.0% of blanks were filtered from the fungal ITS dataset. Most blanks were subsequently removed during demultiplexing, i.e., they had zero sequence reads. The few blanks that remained after filtering resembled other deeply sequenced samples, suggesting a low rate of sporadic cross-contamination. Given the very low number of blank samples that passed demultiplexing, denoising, and filtering, and the multiple filtering methods that were employed on both biological and blank samples, we anticipate that a very low rate of contamination remains in the biological samples. After all filtering steps, a total of 22,201,343 bacterial V4 sequences remained in the dataset (representing 41,588 ASVs) with a median count of 10,095 sequences per sample; and a total of 24,594,392 fungal ITS sequences (representing 16,871 ASVs) with a median count of 11,141 sequences per sample.

QIIME 2’s q2-diversity plugin was used to calculate microbiome beta diversity (between-sample diversity) with ASVs using weighted and unweighted UniFrac distance (35), and with ASVs and k-mer frequencies using Bray-Curtis dissimilarity (88) and Jaccard distance (37), and to perform principal coordinate analysis (PCoA) on ASV-based diversity metrics. The q2-diversity plugin was also used to calculate alpha diversity (within-sample diversity), and q2-longitudinal (89) was used to perform ANOVA tests. Feature tables were evenly subsampled at 1,000 sequences per sample prior to diversity analyses. PCoA ordination plots were built using matplotlib (90) from principal coordinate ordinations calculated by q2-diversity. Two-way permutational multivariate analysis of variance (PERMANOVA) tests (39) (as implemented in the adonis method in the vegan R package (91), wrapped via the q2-diversity plugin) were performed to test whether beta diversity estimates were significantly associated with indoor environmental conditions: both daily mean and weekly coefficient of variation (CV%) of temperature, relative humidity, and CO_2_. For k-mer-based diversity metrics, PERMANOVA and pairwise PERMANOVA tests were implemented in R using the packages vegan and pairwiseAdonis, respectively. Here, PERMANOVA tests focused on testing whether there were significant differences in microbial communities depending broadly on sample type or facility, and pairwise PERMANOVA tests focused on testing whether there were significant differences in the microbial communities between select sample types (raw materials, fermented food products, processing equipment, and building surfaces) within facilities of the same product (wine or cheese).

Canonical correspondence analysis (41) (as implemented in scikit-bio (http://scikit-bio.org/)) was also used as an exploratory method to discover associations between samples, microbial features (as response variables), and these same indoor conditions (as explanatory variables). Random Forest machine-learning regression models (45) were used (as implemented in the q2-sample-classifier plugin (46)), to examine associations between indoor environmental conditions (as target values) and microbial features (as predictive features). Predictive features were assembled from the combination of both fungal and bacterial ASVs and taxonomic abundances at phylum through species levels. Forests were grown with 300 trees and used to predict target values for each sample using a 10-fold cross-validation scheme (i.e., 10 predictive models were trained using 9/10 of the samples; all samples were present in the training sets 9 times, and in the test set once). Multinomial regression reference frames implemented in the songbird python package (42) were used to test differential rank abundance of ASVs as a function of each environmental measurement.

### Remote sensor data

Temperature, relative humidity, carbon dioxide, and volatile organic compound concentrations were monitored continuously using a wireless sensor system as described previously (40).

## Acknowledgments

This research was funded in part by funding from the Alfred P.Sloan Foundation to the Biology and the Built Environment Center (to KLB), the Peter J. Shields Endowed Chair in Dairy Food Science (to DAM), the Stephen Sinclair Scott Endowed Chair in Enology (to RBB), and ETH Zurich (to NAB). The authors would like to thank the food producers for their invaluable cooperation in this study, giving us access to and information about their facilities.

