## Supplemental figures and tables for "Indoor environmental conditions shape the microbial landscape of food production facilities across space and time"

**Table S1.** Types of swabbed/sampled materials for each sample type in creameries.

| Sample Type | Material | Site | Count | Count (sequenced) |
|---|---|---|---|---|
| Building air | Air | Air production | 11 | 3 |
| Building surface swab | Ceiling nonproduction | Ceiling nonproduction | 12 | 6 |
| | Ceiling production | Ceiling production | 27 | 15 |
| | Floor nonproduction | Floor nonproduction | 60 | 54 |
| | Floor production | Floor production | 84 | 77 |
| | Milk container | Bulk tank | 1 | 0 |
| | Packaging equipment | Surface nonproduction | 6 | 4 |
| | Surface nonproduction | Surface nonproduction | 89 | 71 |
| | Surface production | Surface production | 49 | 31 |
| Building system | Drain | Drain | 27 | 22 |
| | HVAC | Air return production | 17 | 13 |
| | | Air supply nonproduction | 4 | 3 |
| | | Air supply production | 9 | 2 |
| | Water supply | Water supply | 11 | 10 |
| Fermented food product | Cheese | Cheese core | 20 | 20 |
| | | Cheese surface | 48 | 47 |
| | Curd | Curd | 5 | 5 |
| | Whey | Whey | 6 | 5 |
| Outdoor air | Air | Air outdoor | 4 | 1 |
| Processing equipment | Brine tank | Brine tank | 6 | 5 |
| | Cheese processing surface | Aging rack | 19 | 16 |
| | | Draining table | 16 | 11 |
| | | Drying rack | 4 | 3 |
| | Cheese utensil | Cheese brush | 5 | 3 |
| | | Cheese mold | 10 | 8 |
| | | Cheese screen | 5 | 4 |
| | | Squeegee | 4 | 3 |
| | Cheese vat | Cheese vat | 5 | 2 |
| | Curd equipment | Curd knives | 9 | 7 |
| | | Curd tank | 8 | 3 |
| | Fluid milk | Raw milk | 2 | 2 |
| | Milk container | Bulk tank | 2 | 0 |
| | | Dipper cup | 2 | 1 |
| | | Milk can | 3 | 0 |
| | Packaging equipment | Condensation trap | 4 | 3 |
| | | Packaging table | 8 | 5 |
| | | Scale | 2 | 2 |
| | | Wrapping paper | 5 | 4 |
| | Pasteurizer | Pasteurizer | 6 | 4 |
| | | Pasteurizer agitator | 4 | 3 |
| | Raw milk | Raw milk hose | 1 | 0 |
| | | Raw milk tank | 1 | 1 |
| | Surface nonproduction | Salt scoop | 3 | 2 |
| | Washing table | Washing table | 2 | 1 |
| | Undefined equipment/utensil | Undefined equipment/utensil | 4 | 4 |
| Raw materials | Brine | Brine | 15 | 15 |
| | Cream | Cream | 6 | 5 |
| | Culture | Culture cheese | 29 | 28 |

| | | | | |
|---|---|---|---|---|
| | Fluid milk | Fluid milk | 8 | 8 |
| | | Fluid milk organic | 1 | 1 |
| | | Raw Milk | 6 | 4 |
| | Plant ingredient | Plant ingredient | 3 | 3 |
| | Rennet | Rennet | 8 | 5 |
| | Salt | Calcium chloride | 2 | 2 |
| | | Sodium chloride | 3 | 2 |
| Vectors | Boots | Boots | 8 | 7 |
| | Cheese cart | Wheel | 23 | 19 |
| | Cleaning equipment | COP tank | 6 | 4 |
| | | Cleaning equipment | 16 | 16 |
| | Undefined vector | Undefined vector | 2 | 2 |
| Waste | Waste | River water | 1 | 1 |

**Table S2.** Types of swabbed/sampled materials for each sample type in wineries.

| Sample Type | Material | Site | Count | Count (sequenced) |
|---|---|---|---|---|
| Building air | Air | Air nonproduction | 14 | 3 |
| | | Air production | 41 | 7 |
| Building surface swab | Ceiling production | Ceiling production | 19 | 13 |
| | Crush equipment | Auger | 4 | 0 |
| | Floor nonproduction | Floor nonproduction | 122 | 53 |
| | Floor production | Floor production | 626 | 274 |
| | Insect | Insect | 1 | 0 |
| | Sink | Sink | 7 | 0 |
| | Surface nonproduction | Surface nonproduction | 244 | 91 |
| | Surface production | Surface production | 251 | 103 |
| | Undefined fermentation hall swab | Undefined fermentation hall swab | 1 | 0 |
| Building system | Drain | Drain | 132 | 59 |
| | HVAC | Humidifier | 13 | 8 |
| | | Air return nonproduction | 44 | 17 |
| | | Air return production | 55 | 10 |
| | | Air supply nonproduction | 19 | 12 |
| | | Air supply production | 50 | 20 |
| | Waste | Wastewater pump | 1 | 0 |
| | | wastewater pond | 17 | 13 |
| Fermented food product | Fermenting juice | Fermenting juice | 35 | 23 |
| | Wine | Wine in barrel | 16 | 9 |
| | | Wine in tank | 8 | 4 |
| Outdoor air | Air | Air outdoor | 11 | 3 |
| Processing equipment | Barrel | Barrel exterior | 43 | 1 |
| | | Barrel exterior new | 3 | 0 |
| | | Barrel interior | 20 | 0 |
| | | Barrel port | 32 | 16 |
| | Capsule | Capsule | 6 | 0 |
| | Crush equipment | Auger | 2 | 0 |
| | | Chute | 12 | 11 |
| | | Conveyer | 7 | 0 |
| | | Destemmer | 33 | 12 |
| | | Grape elevator | 53 | 33 |
| | | Grape hopper | 29 | 12 |

| | | | | |
|---|---|---|---|---|
| | | Juice pan | 4 | 0 |
| | | Must line | 61 | 35 |
| | | Skin pan | 3 | 0 |
| | | Sorting table | 36 | 7 |
| | Fermenter | Fermenter exterior wall | 51 | 28 |
| | | Fermenter inner wall | 289 | 64 |
| | | Fermenter port | 475 | 143 |
| | Grape bin | Grape bin | 13 | 5 |
| | Heat exchanger | Heat exchanger | 3 | 3 |
| | Hose | Juice hose | 10 | 5 |
| | | Wine hose | 70 | 11 |
| | Packaging equipment | Bottle cap | 1 | 0 |
| | | Bottle empty | 1 | 0 |
| | | Bottler bowl | 8 | 0 |
| | | Filler | 4 | 0 |
| | | Wine filter | 3 | 0 |
| | Pour spout | Pour spout | 8 | 0 |
| | Press | Basket press | 7 | 6 |
| | | Juice pan | 23 | 0 |
| | | Press inlet | 9 | 5 |
| | | Press outlet | 30 | 15 |
| | | Press tube | 22 | 12 |
| | | Skin pan | 13 | 0 |
| | Pump | Juice pump | 4 | 3 |
| | | Must pump | 16 | 0 |
| | | Pump | 49 | 24 |
| | Racking arm | Racking arm | 3 | 1 |
| | Surface production | Surface production | 8 | 0 |
| | Tank agitator | Tank agitator | 7 | 0 |
| | Processing line | Processing line | 2 | 0 |
| Raw materials | Culture | Culture yeast | 4 | 3 |
| | Grape | Grape | 23 | 15 |
| | Juice | Juice | 29 | 20 |
| | Must | Must | 19 | 16 |
| Soil | Soil | Soil | 5 | 4 |
| Vectors | Boots | Boots | 107 | 28 |
| | Cleaning equipment | Cleaning equipment | 18 | 12 |
| | Forklift | Wheel | 14 | 8 |
| | Insect | Insect | 33 | 19 |
| | Rag | Rag | 4 | 0 |
| Waste | Waste | Grape waste | 5 | 2 |
| | | Pomace | 11 | 7 |
| | | Rain water | 1 | 1 |
| | | Waste bin | 4 | 3 |
| | | Wastewater | 1 | 0 |

**Table S3**. PERMANOVA analysis of creamery and winery 16S rRNA gene and fungal ITS data from k-mer-based Bray Curtis dissimilarity and Jaccard distance metrics.

| Facility Type | Target | Metric | Variable | Df | $R^2$ | P-value |
|---|---|---|---|---|---|---|

| | | | | | | |
|---|---|---|---|---|---|---|
| Creameries | Bacteria | Bray-Curtis | Facility | 1 | 0.048 | 0.001 |
| | | | Sample Type | 6 | 0.144 | 0.001 |
| | | Jaccard | Facility | 1 | 0.024 | 0.001 |
| | | | Sample Type | 6 | 0.117 | 0.001 |
| | Fungi | Bray-Curtis | Facility | 1 | 0.128 | 0.001 |
| | | | Sample Type | 6 | 0.110 | 0.001 |
| | | Jaccard | Facility | 1 | 0.045 | 0.001 |
| | | | Sample Type | 6 | 0.090 | 0.001 |
| Wineries | Bacteria | Bray-Curtis | Facility | 2 | 0.029 | 0.001 |
| | | | Sample Type | 7 | 0.055 | 0.001 |
| | | Jaccard | Facility | 2 | 0.027 | 0.001 |
| | | | Sample Type | 7 | 0.053 | 0.001 |
| | Fungi | Bray-Curtis | Facility | 2 | 0.031 | 0.001 |
| | | | Sample Type | 7 | 0.064 | 0.001 |
| | | Jaccard | Facility | 2 | 0.034 | 0.001 |
| | | | Sample Type | 7 | 0.059 | 0.001 |

**Table S4.** Pairwise PERMANOVA tests between select sample types - non-k-mer-based metrics.

| Facility Type | Target | Metric* | Group 1 | Group 2 | N | pF* | p-value | q-value |
|---|---|---|---|---|---|---|---|---|
| Creameries | Bacteria | Bray-Curtis | Building surface swab | Fermented food product | 269 | 30.19 | 0.001 | 0.001 |
| | | | | Processing equipment | 297 | 6.90 | 0.001 | 0.001 |
| | | | | Raw materials | 279 | 13.71 | 0.001 | 0.001 |
| | | | Fermented food product | Processing equipment | 138 | 11.81 | 0.001 | 0.001 |
| | | | | Raw materials | 120 | 7.93 | 0.001 | 0.001 |
| | | | Processing equipment | Raw materials | 148 | 4.07 | 0.003 | 0.003 |
| | | Jaccard | Building surface swab | Fermented food product | 269 | 11.78 | 0.001 | 0.001 |
| | | | | Processing equipment | 297 | 4.82 | 0.001 | 0.001 |
| | | | | Raw materials | 279 | 10.21 | 0.001 | 0.001 |
| | | | Fermented food product | Processing equipment | 138 | 3.78 | 0.001 | 0.001 |
| | | | | Raw materials | 120 | 2.50 | 0.002 | 0.002 |
| | | | Processing equipment | Raw materials | 148 | 3.81 | 0.001 | 0.001 |
| | | uUniFrac | Building surface swab | Fermented food product | 269 | 23.47 | 0.001 | 0.001 |
| | | | | Processing equipment | 297 | 9.49 | 0.001 | 0.001 |
| | | | | Raw materials | 279 | 23.07 | 0.001 | 0.001 |
| | | | Fermented food product | Processing equipment | 138 | 6.11 | 0.001 | 0.001 |
| | | | | Raw materials | 120 | 1.66 | 0.074 | 0.076 |
| | | | Processing equipment | Raw materials | 148 | 6.87 | 0.001 | 0.001 |
| | | wUniFrac | Building surface swab | Fermented food product | 269 | 61.47 | 0.001 | 0.001 |

| | | | | | | | | |
|---|---|---|---|---|---|---|---|---|
| | | | | Processing equipment | 297 | 11.60 | 0.001 | 0.001 |
| | | | | Raw materials | 279 | 39.21 | 0.001 | 0.001 |
| | | | Fermented food product | Processing equipment | 138 | 16.30 | 0.001 | 0.001 |
| | | | | Raw materials | 120 | 4.45 | 0.008 | 0.009 |
| | | | Processing equipment | Raw materials | 148 | 7.40 | 0.002 | 0.002 |
| | Fungi | Bray-Curtis | Building surface swab | Fermented food product | 262 | 17.43 | 0.001 | 0.001 |
| | | | | Processing equipment | 268 | 2.50 | 0.038 | 0.040 |
| | | | | Raw materials | 240 | 18.25 | 0.001 | 0.001 |
| | | | Fermented food product | Processing equipment | 146 | 7.56 | 0.002 | 0.002 |
| | | | | Raw materials | 118 | 25.77 | 0.001 | 0.001 |
| | | | Processing equipment | Raw materials | 124 | 11.45 | 0.001 | 0.001 |
| | | Jaccard | Building surface swab | Fermented food product | 262 | 15.96 | 0.001 | 0.001 |
| | | | | Processing equipment | 268 | 6.78 | 0.001 | 0.001 |
| | | | | Raw materials | 240 | 4.74 | 0.001 | 0.001 |
| | | | Fermented food product | Processing equipment | 146 | 4.03 | 0.001 | 0.001 |
| | | | | Raw materials | 118 | 8.99 | 0.001 | 0.001 |
| | | | Processing equipment | Raw materials | 124 | 4.85 | 0.001 | 0.001 |
| Wineries | Bacteria | Bray-Curtis | Building surface swab | Fermented food product | 489 | 7.14 | 0.001 | 0.001 |
| | | | | Processing equipment | 828 | 8.15 | 0.001 | 0.001 |
| | | | | Raw materials | 490 | 6.91 | 0.001 | 0.001 |
| | | | Fermented food product | Processing equipment | 389 | 4.22 | 0.001 | 0.001 |
| | | | | Raw materials | 51 | 2.66 | 0.003 | 0.003 |
| | | | Processing equipment | Raw materials | 390 | 4.53 | 0.001 | 0.001 |
| | | Jaccard | Building surface swab | Fermented food product | 489 | 4.56 | 0.001 | 0.001 |
| | | | | Processing equipment | 828 | 5.49 | 0.001 | 0.001 |
| | | | | Raw materials | 490 | 4.17 | 0.001 | 0.001 |
| | | | Fermented food product | Processing equipment | 389 | 2.90 | 0.001 | 0.001 |
| | | | | Raw materials | 51 | 1.62 | 0.008 | 0.009 |
| | | | Processing equipment | Raw materials | 390 | 2.72 | 0.001 | 0.001 |
| | | uUniFrac | Building surface swab | Fermented food product | 489 | 10.17 | 0.001 | 0.001 |
| | | | | Processing equipment | 828 | 8.38 | 0.001 | 0.001 |
| | | | | Raw materials | 490 | 6.76 | 0.001 | 0.001 |
| | | | Fermented food product | Processing equipment | 389 | 6.38 | 0.001 | 0.001 |
| | | | | Raw materials | 51 | 1.97 | 0.011 | 0.012 |
| | | | Processing equipment | Raw materials | 390 | 3.69 | 0.001 | 0.001 |
| | | wUniFrac | Building surface swab | Fermented food product | 489 | 17.33 | 0.001 | 0.001 |
| | | | | Processing equipment | 828 | 9.76 | 0.001 | 0.001 |
| | | | | Raw materials | 490 | 15.17 | 0.001 | 0.001 |

| | | | | | | | | |
|---|---|---|---|---|---|---|---|---|
| | | | Fermented food product | Processing equipment | 389 | 8.78 | 0.001 | 0.001 |
| | | | | Raw materials | 51 | 1.91 | 0.083 | 0.084 |
| | | | Processing equipment | Raw materials | 390 | 6.73 | 0.001 | 0.001 |
| | Fungi | Bray-Curtis | Building surface swab | Fermented food product | 523 | 11.94 | 0.001 | 0.001 |
| | | | | Processing equipment | 887 | 14.84 | 0.001 | 0.001 |
| | | | | Raw materials | 549 | 19.10 | 0.001 | 0.001 |
| | | | Fermented food product | Processing equipment | 418 | 6.86 | 0.001 | 0.001 |
| | | | | Raw materials | 80 | 1.76 | 0.084 | 0.084 |
| | | | Processing equipment | Raw materials | 444 | 9.53 | 0.001 | 0.001 |
| | | Jaccard | Building surface swab | Fermented food product | 523 | 4.87 | 0.001 | 0.001 |
| | | | | Processing equipment | 887 | 8.37 | 0.001 | 0.001 |
| | | | | Raw materials | 549 | 10.22 | 0.001 | 0.001 |
| | | | Fermented food product | Processing equipment | 418 | 2.88 | 0.001 | 0.001 |
| | | | | Raw materials | 80 | 2.23 | 0.001 | 0.001 |
| | | | Processing equipment | Raw materials | 444 | 6.09 | 0.001 | 0.001 |

* Abbreviations: uUniFrac = Unweighted UniFrac; wUniFrac = Weighted UniFrac; pF = pseudo-F value.

**Table S5.** Pairwise PERMANOVA tests between select sample types - k-mer-based metrics.

| Facility Type | Target | Metric | Group 1 | Group 2 | $R^2$ | q-value |
|---|---|---|---|---|---|---|
| Creameries | Bacteria | Bray-Curtis | Building surface swab | Fermented food product | 0.180 | 0.001 |
| | | | | Processing equipment | 0.041 | 0.001 |
| | | | | Raw materials | 0.110 | 0.001 |
| | | | Fermented food product | Processing equipment | 0.103 | 0.001 |
| | | | | Raw materials | 0.054 | 0.001 |
| | | | Processing equipment | Raw materials | 0.043 | 0.001 |
| | | Jaccard | Building surface swab | Fermented food product | 0.129 | 0.001 |
| | | | | Processing equipment | 0.049 | 0.001 |
| | | | | Raw materials | 0.110 | 0.001 |
| | | | Fermented food product | Processing equipment | 0.049 | 0.001 |
| | | | | Raw materials | 0.014 | 0.097 |
| | | | Processing equipment | Raw materials | 0.041 | 0.001 |
| | Fungi | Bray-Curtis | Building surface swab | Fermented food product | 0.075 | 0.001 |
| | | | | Processing equipment | 0.012 | 0.021 |
| | | | | Raw materials | 0.088 | 0.001 |
| | | | Fermented food product | Processing equipment | 0.051 | 0.002 |
| | | | | Raw materials | 0.196 | 0.001 |
| | | | Processing equipment | Raw materials | 0.096 | 0.001 |
| | | Jaccard | Building surface swab | Fermented food product | 0.084 | 0.001 |
| | | | | Processing equipment | 0.043 | 0.001 |

| | | | | | | |
|---|---|---|---|---|---|---|
| | | | | Raw materials | 0.040 | 0.001 |
| | | | Fermented food product | Processing equipment | 0.020 | 0.016 |
| | | | | Raw materials | 0.101 | 0.001 |
| | | | Processing equipment | Raw materials | 0.057 | 0.001 |
| Wineries | Bacteria | Bray-Curtis | Building surface swab | Fermented food product | 0.040 | 0.001 |
| | | | | Processing equipment | 0.015 | 0.001 |
| | | | | Raw materials | 0.031 | 0.001 |
| | | | Fermented food product | Processing equipment | 0.029 | 0.001 |
| | | | | Raw materials | 0.057 | 0.004 |
| | | | Processing equipment | Raw materials | 0.019 | 0.001 |
| | | Jaccard | Building surface swab | Fermented food product | 0.055 | 0.001 |
| | | | | Processing equipment | 0.014 | 0.001 |
| | | | | Raw materials | 0.036 | 0.001 |
| | | | Fermented food product | Processing equipment | 0.037 | 0.001 |
| | | | | Raw materials | 0.035 | 0.028 |
| | | | Processing equipment | Raw materials | 0.020 | 0.001 |
| | Fungi | Bray-Curtis | Building surface swab | Fermented food product | 0.052 | 0.001 |
| | | | | Processing equipment | 0.019 | 0.001 |
| | | | | Raw materials | 0.055 | 0.001 |
| | | | Fermented food product | Processing equipment | 0.037 | 0.001 |
| | | | | Raw materials | 0.030 | 0.056 |
| | | | Processing equipment | Raw materials | 0.033 | 0.001 |
| | | Jaccard | Building surface swab | Fermented food product | 0.036 | 0.001 |
| | | | | Processing equipment | 0.016 | 0.001 |
| | | | | Raw materials | 0.048 | 0.001 |
| | | | Fermented food product | Processing equipment | 0.027 | 0.001 |
| | | | | Raw materials | 0.036 | 0.016 |
| | | | Processing equipment | Raw materials | 0.035 | 0.001 |

**Table S6.** Analysis of variance of observed ASVs (richness) of bacteria and fungi in creameries and wineries.*

| | Creamery Bacteria | | Winery Bacteria | | Creamery Fungi | | Winery Fungi | |
|---|---|---|---|---|---|---|---|---|
| | Estimate | P-value | Estimate | P-value | Estimate | P-value | Estimate | P-value |
| Facility | | 0.078 | | 0.114 | | 0.100 | | 0.258 |
| Room Type | | 0.473 | | 0.148 | | 0.923 | | 0.052 |
| Sample Type | | 0.022 | | 0.000 | | 0.287 | | 0.000 |
| Daily Mean Temp | -9.44 | 0.017 | -66.11 | 0.007 | 1.26 | 0.558 | -4.35 | 0.735 |
| Daily Mean RH | -2.71 | 0.038 | -21.57 | 0.003 | 0.61 | 0.388 | -5.65 | 0.117 |

| | | | | | | | | |
|---|---|---|---|---|---|---|---|---|
| Daily Mean CO2 | 0.00 | 0.770 | 0.00 | 0.949 | 0.00 | 0.947 | -0.02 | 0.001 |
| Weekly CV Temp | 374.47 | 0.378 | 10300.00 | 0.040 | 40.93 | 0.862 | 1832.82 | 0.491 |
| Weekly CV RH | -242.10 | 0.281 | -906.94 | 0.532 | 58.60 | 0.631 | -1701.73 | 0.005 |
| Weekly CV CO2 | 83.85 | 0.109 | 519.23 | 0.098 | -8.92 | 0.804 | -24.72 | 0.882 |

* Observed richness was estimated on feature tables evenly sampled at 1000 sequences per sample. Ordinary least squares regression coefficient estimates are provided for continuous variables. These estimates indicate the direction of effect. Estimates are not provided for multiclass categorical variables.

**Table S7.** Random Forest regressor predictive accuracy of indoor environment based on microbiome measurements.

| Facility | Variable | MSE | $R^2$ | P-value | Slope |
|---|---|---|---|---|---|
| Winery | Daily Mean Temp | 6.98 | 0.56 | < 0.001 | 0.51 |
| | Daily Mean Humidity | 60.38 | 0.50 | < 0.001 | 0.46 |
| | Daily Mean CO2 | 1949269.84 | 0.41 | < 0.001 | 0.37 |
| | Weekly CV Temp | 0.00 | 0.04 | 0.005 | 0.07 |
| | Weekly CV Humidity | 0.00 | 0.60 | < 0.001 | 0.44 |
| | Weekly Mean CO2 | 2161834.80 | 0.43 | < 0.001 | 0.38 |
| Creamery | Daily Mean Temp | 9.91 | 0.21 | < 0.001 | 0.22 |
| | Daily Mean Humidity | 79.37 | 0.19 | < 0.001 | 0.22 |
| | Daily Mean CO2 | 172312.09 | 0.18 | < 0.001 | 0.15 |
| | Weekly CV Temp | 0.00 | 0.11 | 0.006 | 0.12 |
| | Weekly CV Humidity | 0.00 | 0.32 | < 0.001 | 0.31 |
| | Weekly Mean CO2 | 197481.97 | 0.15 | 0.001 | 0.15 |

* Abbreviations: V4 = 16S rRNA gene V4 domain; MSE = mean squared error; CV = coefficient of variation.

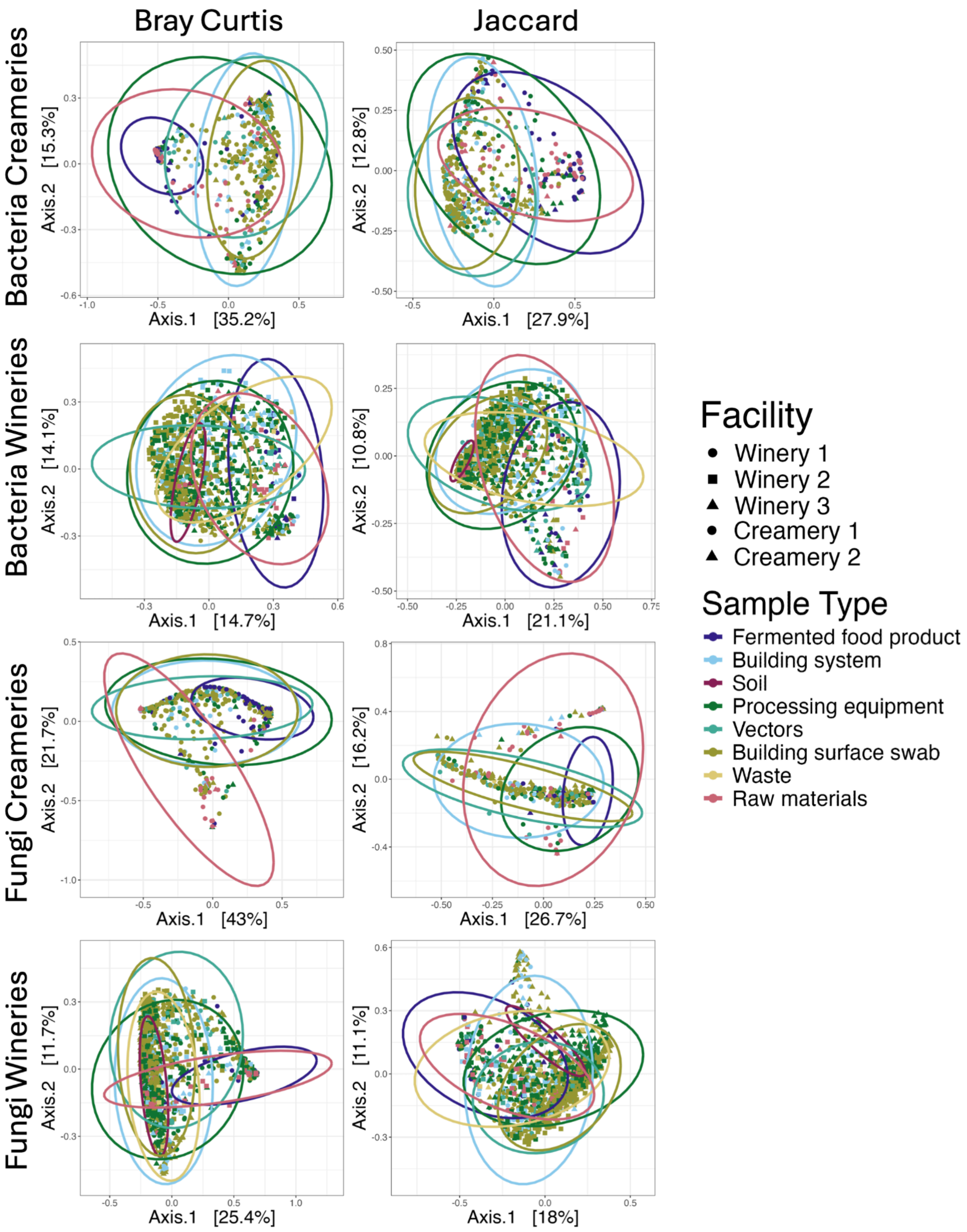


**Figure S1. Kmer-based principal coordinate analysis of winery and creamery samples**. Principal coordinate analysis visualization of kmer-based beta diversity metrics for both bacteria (V4) and fungi (BITS) evaluated with Bray-Curtis and Jaccard. Samples (points) are color-coded by sample type and shape represents different facilities. Ellipses represent the 95% confidence level for the centroid of each sample type, based on a multivariate t-distribution.

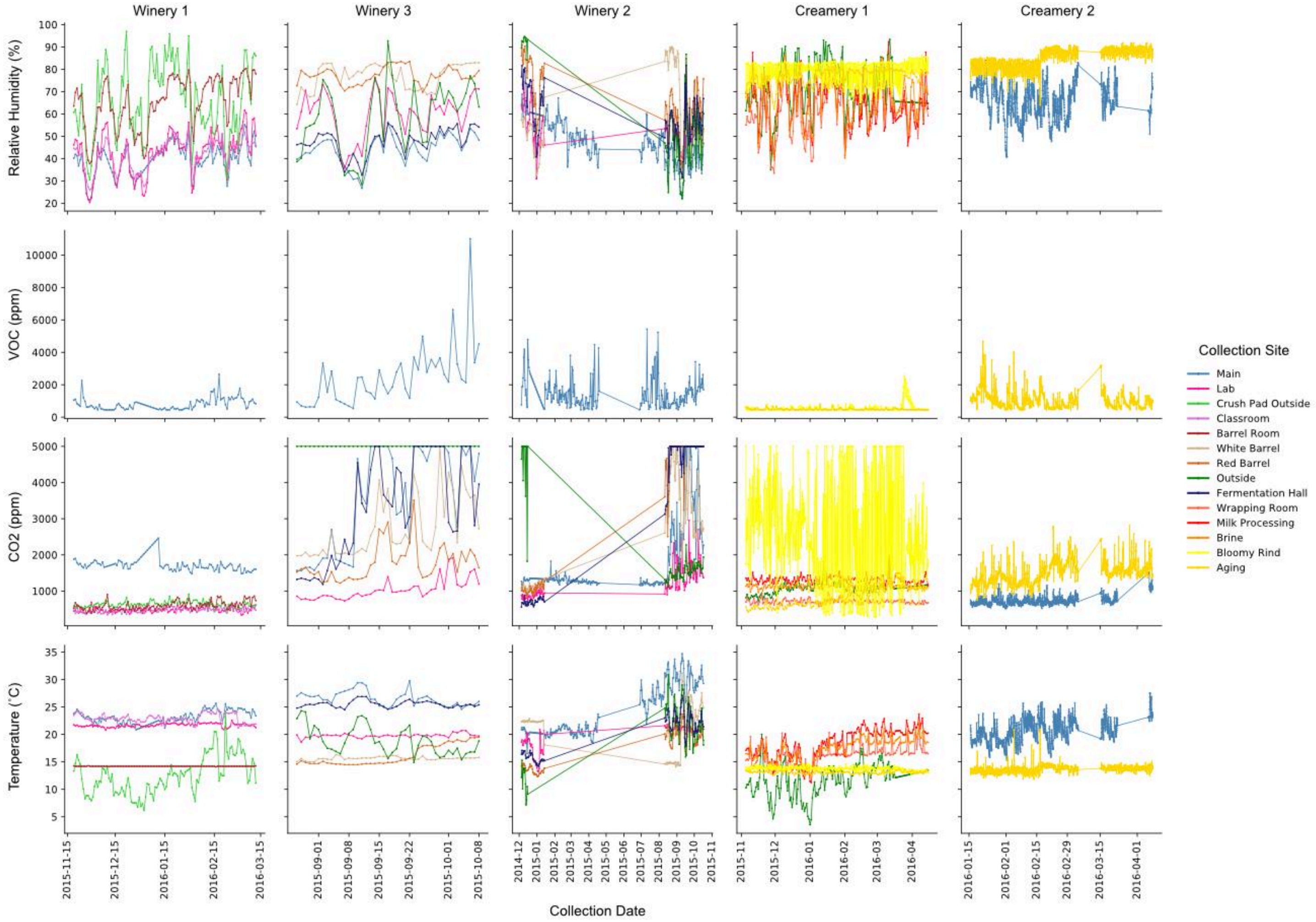


**Figure S2. Indoor environmental monitoring indicates different degrees of volatility.** Line plots display the daily average relative humidity, VOC (ppm), $CO_2$ (ppm), and temperature (˚C) of each remote sensor, color-coded by site function, in each food processing facility. Note that periods of linearity were caused by lapses in sensor readings. The $CO_2$ measurements are clipped due to the upper detection limit of the sensor.

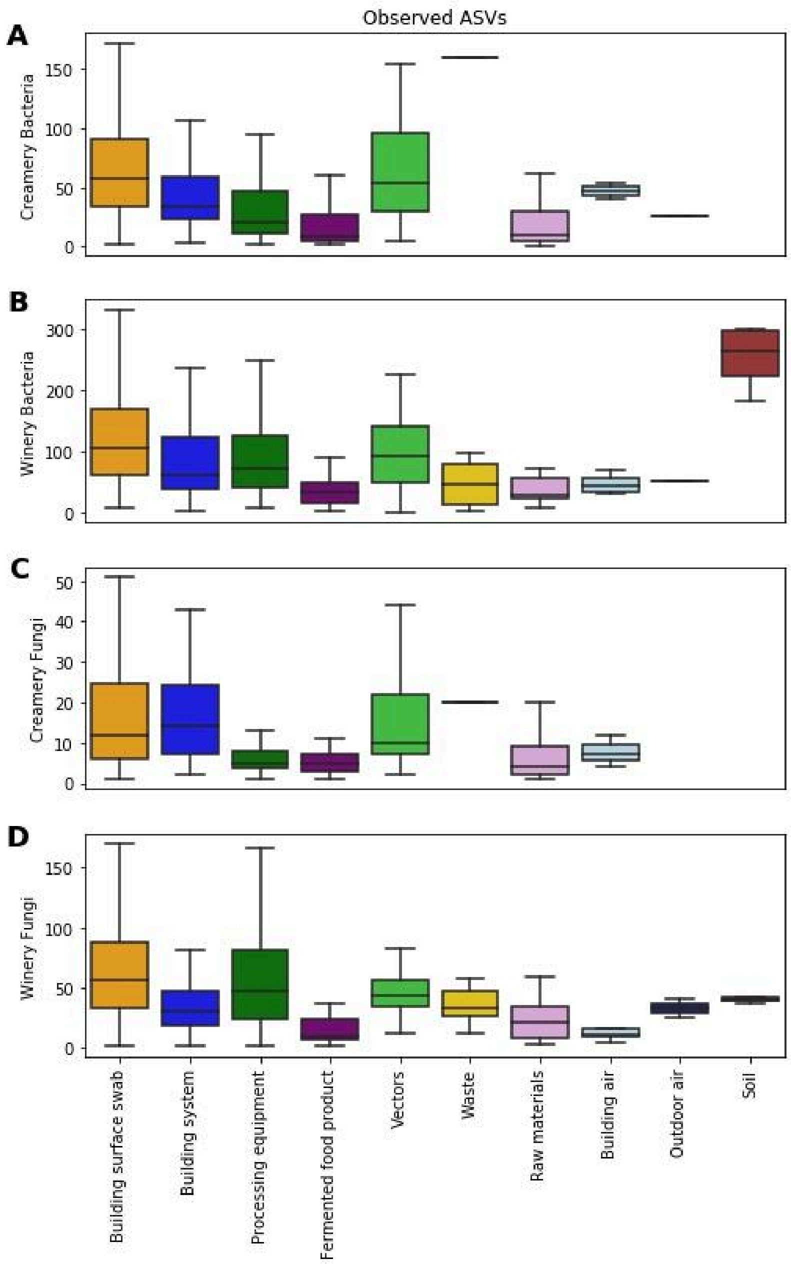


**Figure S3. Bacterial and fungal richness (observed ASVs) in each sample type.** Boxplots display quartile richness estimates of bacteria (panes A and B) and fungi (panels C and D) for all sample types averaged across creameries (panels A and C) and wineries (panels B and D). To normalize for uneven sampling depths, estimates were calculated on feature tables that were evenly sampled at 1000 sequences per sample. In general, differences in sample type were driven by significantly lower richnesses in raw materials and fermented food products than in building surfaces. Processing equipment also exhibited significantly lower bacterial and fungal richness estimates than non-processing surfaces ($P < 0.05$), but still higher than raw materials and fermented food products, with the exception of fungal richness in creameries.

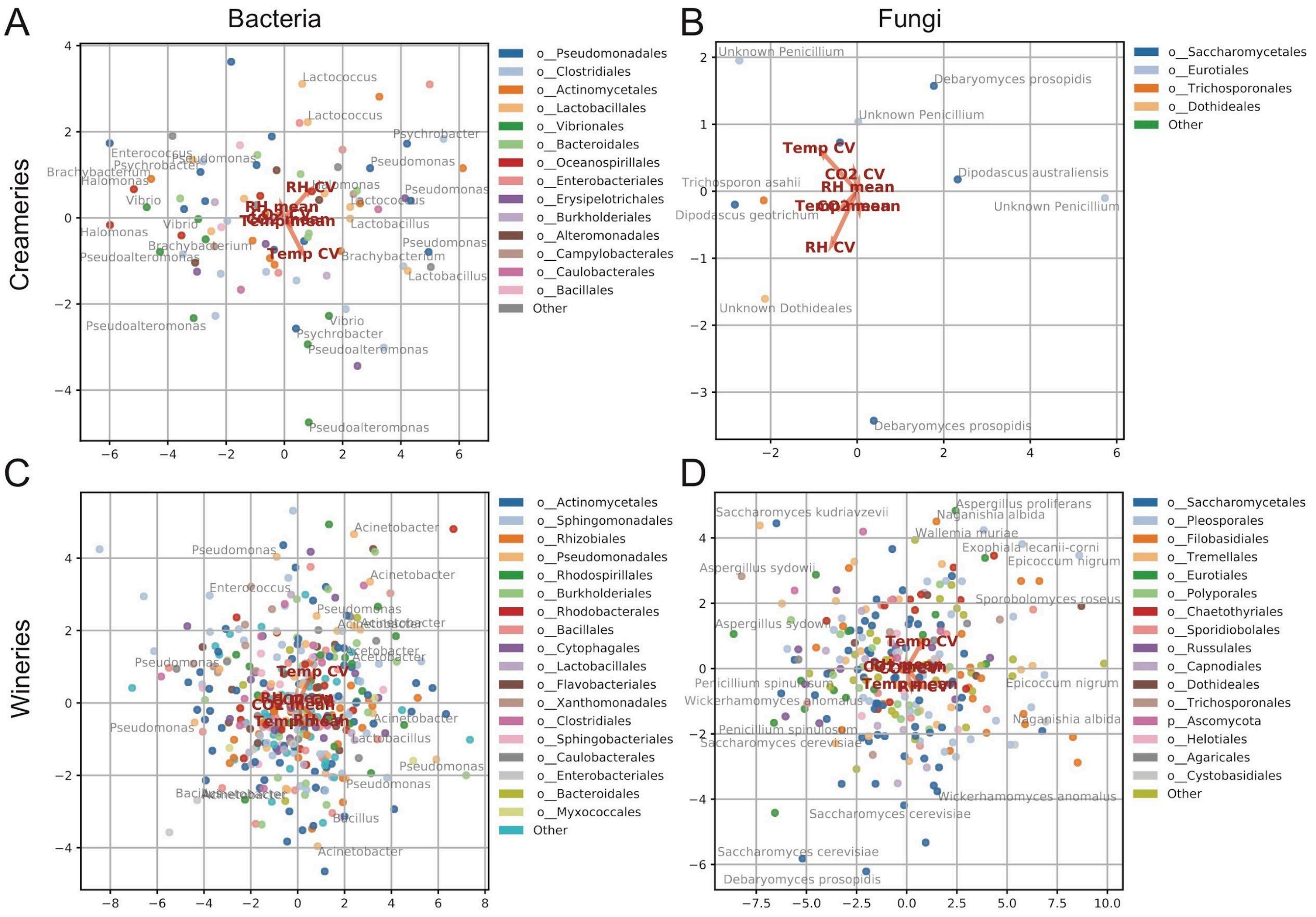


**Figure S4. Multinomial regression differentials biplots indicate associations between indoor environment and microbial community composition.** Multinomial regression of daily mean temperature (Temp mean), humidity (RH mean), and $CO_2$ ($CO_2$ mean) and weekly coefficient of variance for temperature (Temp CV), humidity (RH CV), and $CO_2$ ($CO_2$ CV) as explanatory variables; and bacterial (left panels) or fungal community compositions (right panels) in creameries (top panels) or wineries (bottom panels) as observations. Each panel displays the co-ordination of variables along axes 1 (horizontal) and 2 (vertical). Only select ASVs with strong associations are labeled (via species label) in each panel. Points (ASVs) in each plot are color-categorized by order or the next-lowest rank if that ASV is incompletely classified; taxon prefixes indicate whether the taxonomic affiliation shown is phylum (p__) or order (o__).

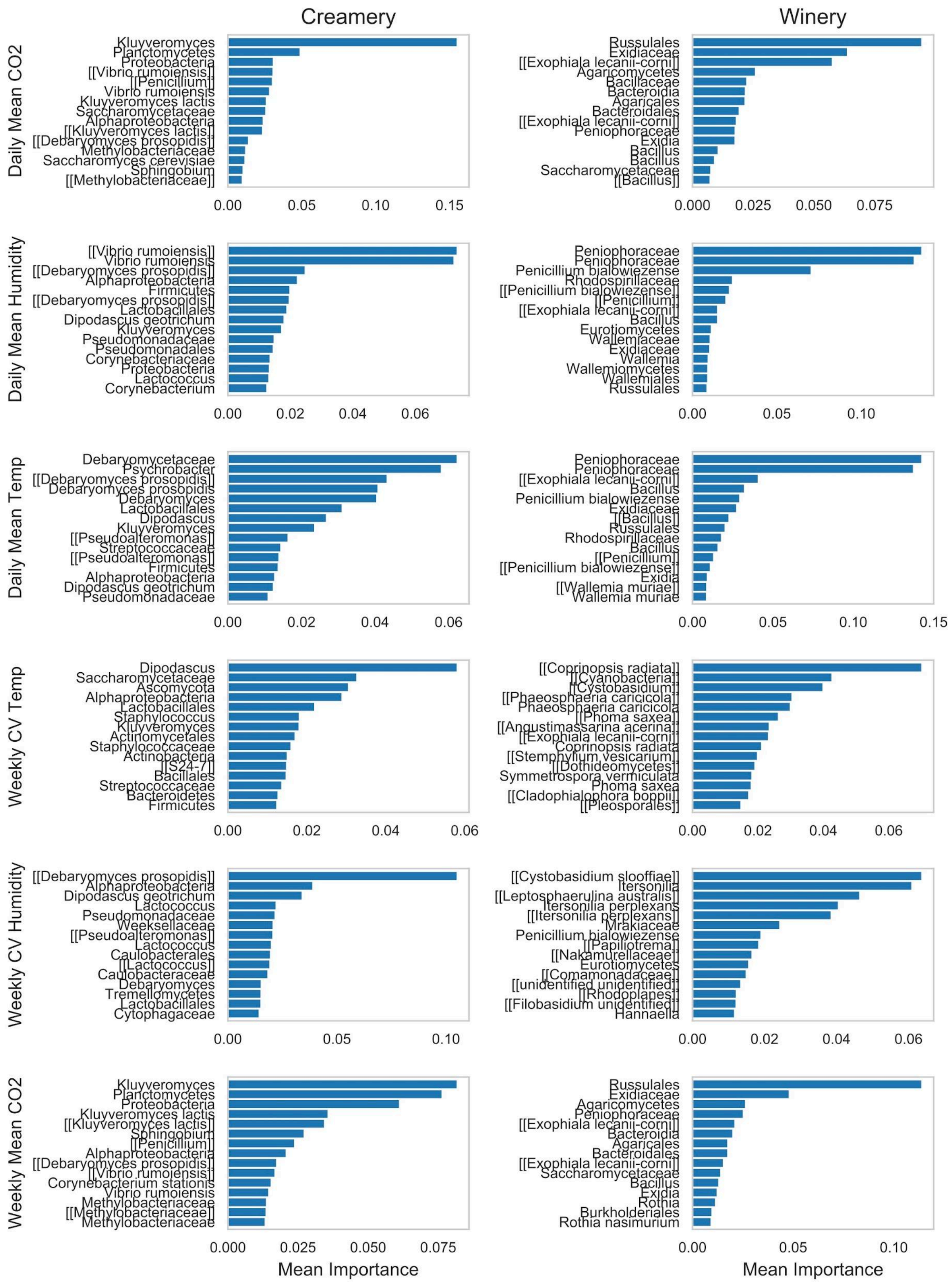


**Figure S5. Random Forest regressors identify bacteria and fungi associated with indoor environmental conditions**. Genus and species name (if identified) of the top fifteen predictive features for each indoor environmental measurement identified by random forest regressor results shown in Table 5. Taxon names displayed in double brackets (e.g., “[[Penicillium]]”) represent ASVs used as predictive features, whereas taxon names without brackets represent the combined abundance of all ASVs assigned that taxonomic affiliation used as predictive features.
